# From spark to wildfire: illuminating single-cell origins of virus infection spread and diversification

**DOI:** 10.1101/2025.10.10.681682

**Authors:** Rahul Ramachandran, Huicheng Shi, John Yin

## Abstract

When a virus encounters a living cell it triggers infection, cell death, and the release of virus progeny. In lab cultures, such progeny spread to near and distant cells, creating a plaque, a visible record of virus growth and cell death. Plaque-based assays have long enabled amplification and quantification of virus particles, but the earliest infections of a plaque remain poorly characterized. Here, we used a recombinant rhinovirus expressing a fluorescent reporter protein to track the start and spread of infection across HeLa cell monolayers at single-cell resolution. We analyzed reporter expression profiles from eight plaques and over 150 cells, reviewing, testing and applying methods of time-series analysis, clustering, and dimensionality reduction. We identified primary infections based on early reporter expression in isolated cells and secondary infections among their neighbors. Primary infected cells exhibited tightly clustered gene expression profiles, while secondary infected cells showed broader heterogeneity in the timing, intensity, and duration of reporter expression, without correspondence to their associated primary cells. This heterogeneity likely reflects a combination of factors, including noisy gene expression, viral genetic changes, and variation in local cellular environments. The rapid diversification of gene expression highlights the power of single-cell imaging to shed light on infection spread.

## INTRODUCTION

The plaque assay is a cornerstone of virology and quantitative biology, widely recognized as a gold standard for measuring the concentration of infectious virus particles. Developed over a century ago, initially to quantify bacteriophage and later to quantify eukaryotic viruses, the assay involves applying a diluted virus sample to a monolayer of susceptible host cells, overlaying the culture with a semi-solid medium such as agar or agarose, and allowing infection cycles to unfold. The resulting plaques —clear zones of lysed or disrupted cells—are counted to calculate the number of infectious virus particles, reported as plaque-forming units (PFU) per milliliter of sample. This quantitative power has made the plaque assay an indispensable tool for characterizing viral infectivity under a variety of experimental conditions [1–3].

Beyond quantifying infectious units, plaques have served as windows into the dynamic interplay between viruses and their hosts. The size, morphology, and growth rate of plaques represent measurable phenotypes that reflect key aspects of viral replication, spread, and host-cell interactions. The plaque growth rate integrates the biophysical coupling of virus diffusion with intracellular amplification, providing a readout of viral fitness in specific host-cell environments [4–6]. Plaque growth rates or plaque size can also be modulated by antiviral compounds, enabling one to test drugs based on their ability to inhibit plaque growth.

Finally, plaques are useful for the clonal amplification of viruses. Plaque formation is initiated by infection of a single cell by a single infectious virus particle, so descendant viruses of the initial infection will be genetically identical (clonal) or homogeneous with respect to their genetics. This virus amplification process, termed plaque purification, has been used to prepare well-defined virus stocks, study the genetic basis of viral traits, or amplify recombinant viruses.

In recent years, technological advances have expanded the utility of plaque assays. Engineered viruses expressing fluorescent reporters, for instance, enable real-time visualization of infection dynamics at single-cell resolution [7,8]. Such innovations have uncovered striking heterogeneity in virus-host interactions [7,9–12], with implications for understanding disease outcomes [12–14]. Yet, the early dynamics of plaque formation—where infection transitions from a single-cell event to multicellular spread— remain poorly understood. Here we use a recombinant rhinovirus engineered to express green fluorescent protein (GFP) to generate dynamic reporter profiles of single-cell infection. Methods to compare such profiles have yet to be established, so we reviewed and tested approaches of time-series analysis, clustering, and dimensionality reduction. Based on initial analyses on toy data sets, we then choose a subset of techniques to analyze our experimental data.

## METHODS

### Plaque growth experiments

Time-series fluorescence data were acquired from HeLa cell monolayers infected with a recombinant human rhinovirus (A16) engineered to express enhanced green fluorescent protein (eGFP) as a reporter of infection during plaque growth. A plaque was initiated when a single infectious virus particle encountered a single cell. Viral gene expression within the infected cell (round-1) led to the accumulation of eGFP, producing detectable fluorescence under an automated fluorescence microscope, as described previously [8]. Upon cell lysis, progeny virus spread to neighboring cells, initiating subsequent rounds of infection. This process resulted in expanding zones of infection, illustrated schematically (Fig. 1a) and captured at single-cell resolution (Fig. 1b).

**Figure 1.**
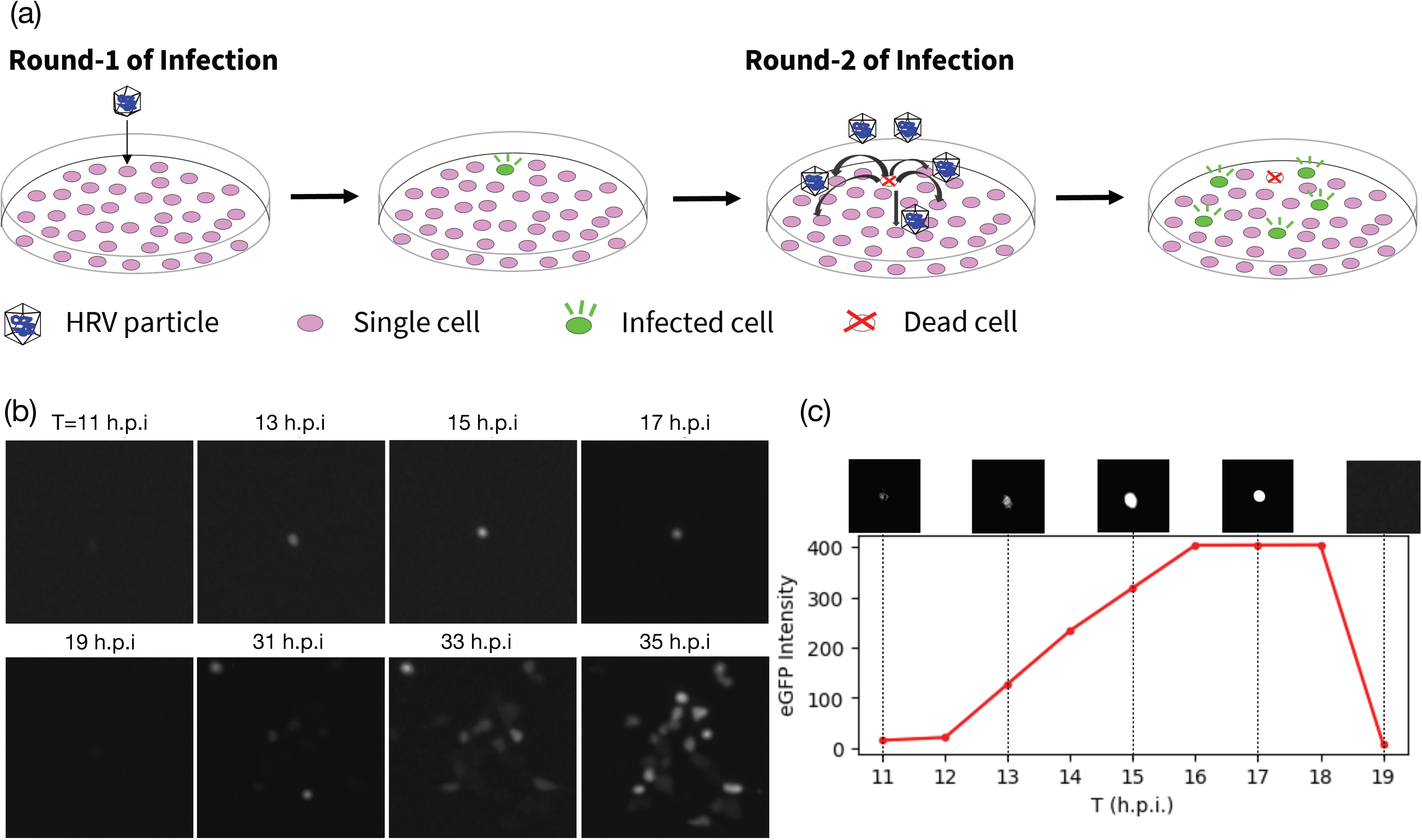
(a) Spread of infection from a single virus-cell encounter, with round-1 representing the primary infection and round-2 the secondary infection. (b)Single-cell resolution fluorescence microscopy images showing spread of infection from a single cell in round-1 to neighboring cells in round-2. (c) Intensity profile of a typical infected cell. Fluorescence intensity increases gradually as more eGFP is produced. Fluorescence intensity drops abruptly as the infection causes the cell to lyse. A fluorescence microscopy image for each data point is shown above the graph.

Infected cultures were imaged at regular intervals to track infection dynamics at single-cell resolution. Images were subjected to background correction and quantitative analysis using JEX software [15], yielding time-series intensity profiles for individual cells in multiple plaques. The typical infection profile for a single cell comprised a gradual rise in fluorescence intensity, reflecting protein accumulation, followed by an abrupt drop coinciding with cell death or lysis (Fig. 1c).

### Data Processing and Cleaning

Intensity profiles were cleaned in three steps (Fig. 2a). (1) Basic cleaning: (a) Only plaques initiated by a single primary infected cell were retained for analysis; plaques arising from multiple primary infections or overlapping plaques were excluded. (b) Missing values (typical sampling ≈ 1 h) were linearly interpolated. (c) On the declining phase, points < 20 a.u. were retained only for the first occurrence; later values were discarded. (2) Recursive smoothing: local minima, attributed to photobleaching or transient perturbations, were iteratively replaced with the mean of neighboring points until eliminated. (3) Artificial tails: if the first or last recorded value exceeded 20 a.u., an additional point was inserted one time step earlier or later, assigned a random value from 0–20 a.u., approximating the microscope’s lower detection limit.

**Figure 2.**
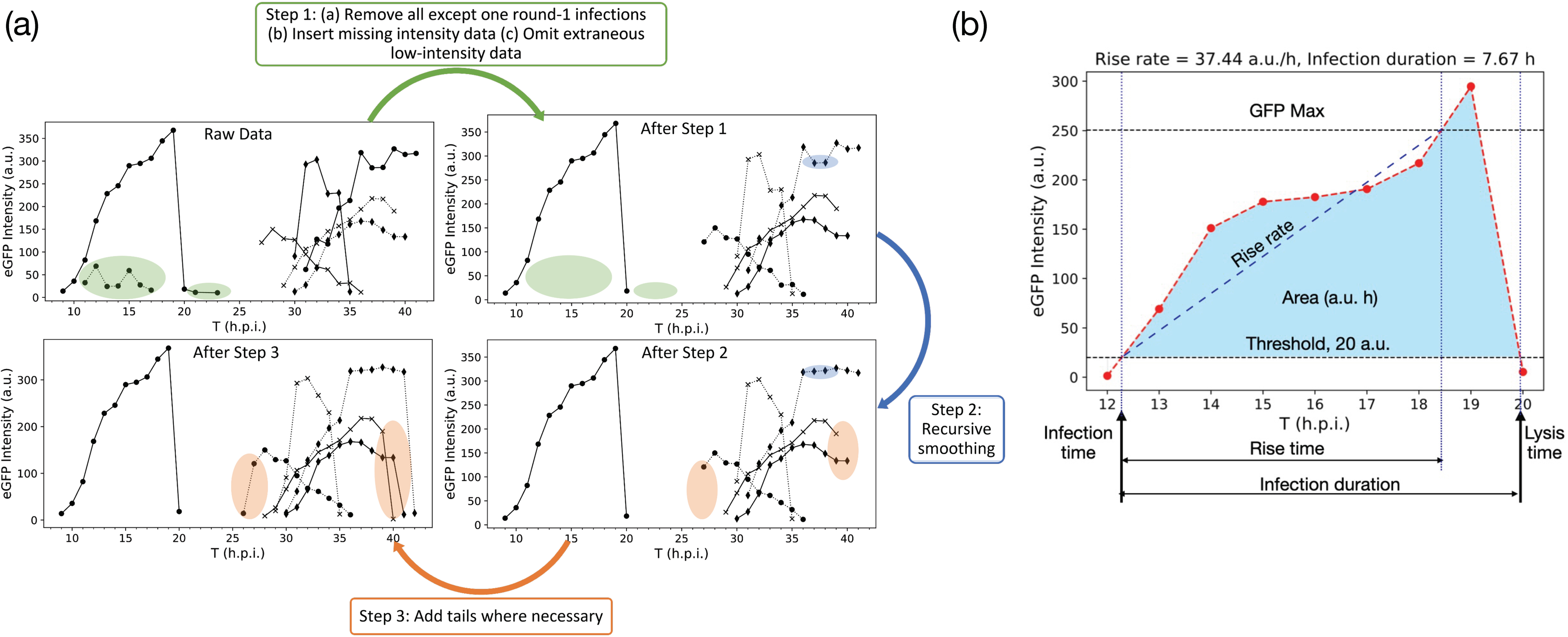
(a) Steps involved in cleaning the time series data. (b) Attributes of a typical intensity profile. Details are presented in the main text.

### Feature Extraction

Different cells exhibit a diversity of fluorescent intensity profiles, and several analytical methods require fixed-length feature vectors. So each single-cell profile was used to estimate its features: rise time, rise rate, infection duration, GFP Max and Area (Fig. 2b). The time series fluorescent data for the expression profile of each infected cell was cleaned as discussed above, and we extracted seven features: infection time (first crossing of 20 a.u.), rise time (interval from infection time to 85 percent of maximum intensity), lysis time (first drop below 20 a.u. after the peak), infection duration (infection time to lysis time), rise rate (slope of the intensity rise), maximum GFP intensity, and the area under the curve between infection and lysis.

These features were standardized by subtracting their mean and dividing by their standard deviation to prevent attributes with larger magnitudes from dominating the analyses.

## Data Analysis

We evaluated a range of dimensionality-reduction, clustering, and sequence-comparison methods to assess their ability to distinguish between primary (round-1) and secondary (round-2) infections and to detect potential plaque-specific or lineage-dependent signatures. The algorithms tested included principal component analysis (PCA) [16–18], kernel PCA [19], DBSCAN [20–23], PCA-DBSCAN [22], K-means clustering [24–26], metric multidimensional scaling (MDS) [27,28], locally linear embedding (LLE) [29–31], t-distributed stochastic neighbor embedding (t-SNE) [32,33], dynamic time warping (DTW) [34–38], and a root mean square error (RMSE) alignment method. Reviews of each algorithm, with literature references and parameter choices, are provided in the Supporting Information 1).

### PCA and kernel PCA

PCA was performed on the standardized data with seven features to project it onto a lower-dimensional sub-space. Seven principal components (PCs) were calculated, and a scree plot was used to visualize the eigenvalues (or variances) explained by each principal component. PCA looked for a linear combination of the attributes that had maximum variance. An attribute was considered important to a PC if the magnitude of its loading score was greater than 1/√*n* where n is the number of samples. The magnitude of an attribute’s loading score indicated the strength of its contribution to the principal component, and its sign indicated whether the influence was positive or negative [17]. Kernel PCA was also performed on the standardized data using a sigmoid kernel, with seven PCs retained for analysis.

### DBSCAN

DBSCAN required two hyperparameters—Eps and MinPts. To determine the optimum value for Eps, we computed the average of the distances of every data point to its k nearest neighbors. The average distances were then plotted in ascending order, and the optimum value was located where the slope of the curve changed abruptly.

Typically, the value of MinPts was chosen based on the heuristic, MinPts ≥ D+1, where D is the dimensionality of the dataset. For large datasets, MinPts was twice the dimensionality. Domain knowledge also needed to be factored in while choosing MinPts. Noisy datasets required a larger value of MinPts. Based on those parameters, DBSCAN classified data as core points, border points, and noise [20].

### K-means

K-means clustering was performed on the standardized dataset with seven features. To determine the optimal number of clusters K for each dataset, both the elbow method [39] and silhouette coefficients [40] were applied. Clustering was conducted over a range of K values, and for each K, the sum of squared distances of samples to their nearest cluster centers (inertia) and the corresponding silhouette coefficients were computed. The silhouette coefficient ranged from +1 to –1, with higher values indicating more distinct and well-separated clusters.

### Metric MDS

Two-dimensional metric MDS was applied to the standardized dataset with seven dimensions. Pairwise Euclidean distances between the samples were used to construct the dissimilarity matrix. A two-dimensional embedding of the dataset was created and optimized such that the pairwise distances between these new embedded points approximated the corresponding values from the dissimilarity matrix closely.

### LLE

LLE was applied to the standardized dataset with seven features. The algorithm reconstructs each data point as a linear combination of its nearest neighbors and computes an embedding that preserves these local relationships in lower dimensions. The number of neighbors, k, was set to eight, chosen to balance preservation of local geometry with robustness against noise. A two-dimensional embedding was generated for visualization and comparison across cells.

### t-SNE

t-SNE was performed on the standardized dataset with seven features to capture nonlinear patterns. The method converts pairwise distances into probabilities that reflect neighborhood similarity and optimizes a two-dimensional map to preserve these relationships. The key hyperparameter, perplexity, which controls the balance between local and global structure, was varied from 5 to 50. This range was chosen to test whether multi-scale organization could be identified in the dataset.

### DTW

The intensity profile of each cell was standardized using the mean and the standard deviation of that cell’s intensities. The distance between the standardized profiles of cell pairs was calculated using the DTW algorithm. If the profiles were similar, then the distance was 0, and if the profiles were dissimilar, then the distance was 1. In most cases, the distance values were between 0 and 1. Once the distance for all possible cell pairs had been calculated, a symmetric distance matrix was constructed.

That matrix was typically visualized using a heat map or dendrogram. For creating the dendrogram, agglomerative hierarchical clustering (HC) was performed with average linkage. In average linkage, the distance between two clusters was the average distance between data points. Metric MDS was also used to visualize the similarity between cells’ profiles across plaques.

### RMSE

We quantified the similarity between two time-series sequences without stretching or compressing them, distinguishing RMSE from DTW. One sequence was shifted relative to the other along the time axis in fixed increments (1 hour, based on the sampling rate), and for each shift, RMSE was computed only over overlapping time points (Fig. 3). This avoided distortions from time-axis scaling while enabling locally meaningful comparisons. For each cell pair, the shift yielding the minimum RMSE was recorded. Repeating this process for all cell pairs produced a symmetric dissimilarity matrix, scaled to range from 0 (identical) to 1 (maximally different). This matrix was visualized as a heat map and a dendrogram to identify potential clusters. Although RMSE could be applied to the first derivative of intensities or to cumulative intensity profiles (one-step growth curves), these transformations amplified noise in low-sampling data. Therefore, all reported results used raw intensity profiles.

**Figure 3.**
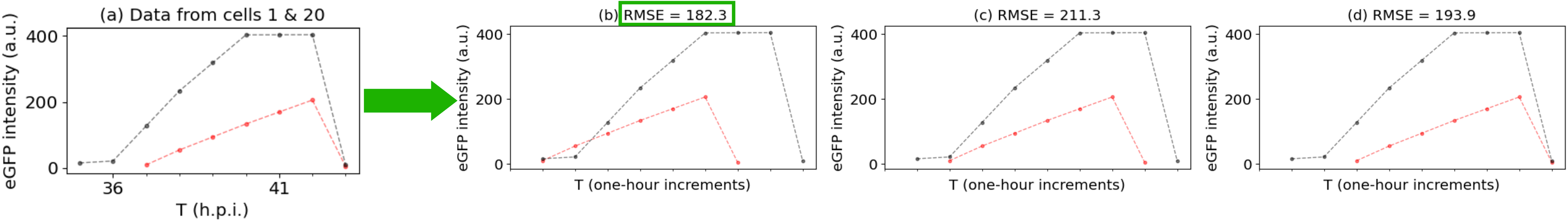
Method for comparing root mean square error (RMSE) between intensities: Frame (a) displays intensity profiles of two cells within a plaque. Subsequently, the profiles are synchronized by shifting one along the time axis in 1-hour increments, and the RMSE between the profiles is computed for each alignment (b-d). The minimum RMSE for that cell pair is documented (c). This process is repeated for all cell pairs considered. Ultimately, a dissimilarity matrix is obtained by combining the minimum RMSEs from all possible pairs.

### Validation on Simulated Data

Before applying these methods to our plaque-growth data, we tested them on synthetic (“toy”) datasets designed to mimic specific features of our data while avoiding experimental noise. These datasets were constructed to explore the effects of environmental variability, genetic variability, and clock-time alignment on clustering performance (Supporting Information 2). Performance on simulated data guided the selection of methods for the experimental analyses.

## RESULTS and DISCUSSION

We analyzed eight plaques comprising 154 single-cell eGFP expression profiles. The number of tracked cells per plaque ranged from 12 to 24, reflecting plaque-to-plaque differences in infection progression and image quality. For each cell, we extracted infection time, lysis time, infection duration, rise rate, maximum GFP intensity, and area under the curve, capturing both temporal and amplitude features of the expression profiles. An overall picture of data was feasible by plotting infection versus lysis times for individual plaques and combining profiles from all cells across all plaques (**Fig. 4**).

**Figure 4.**
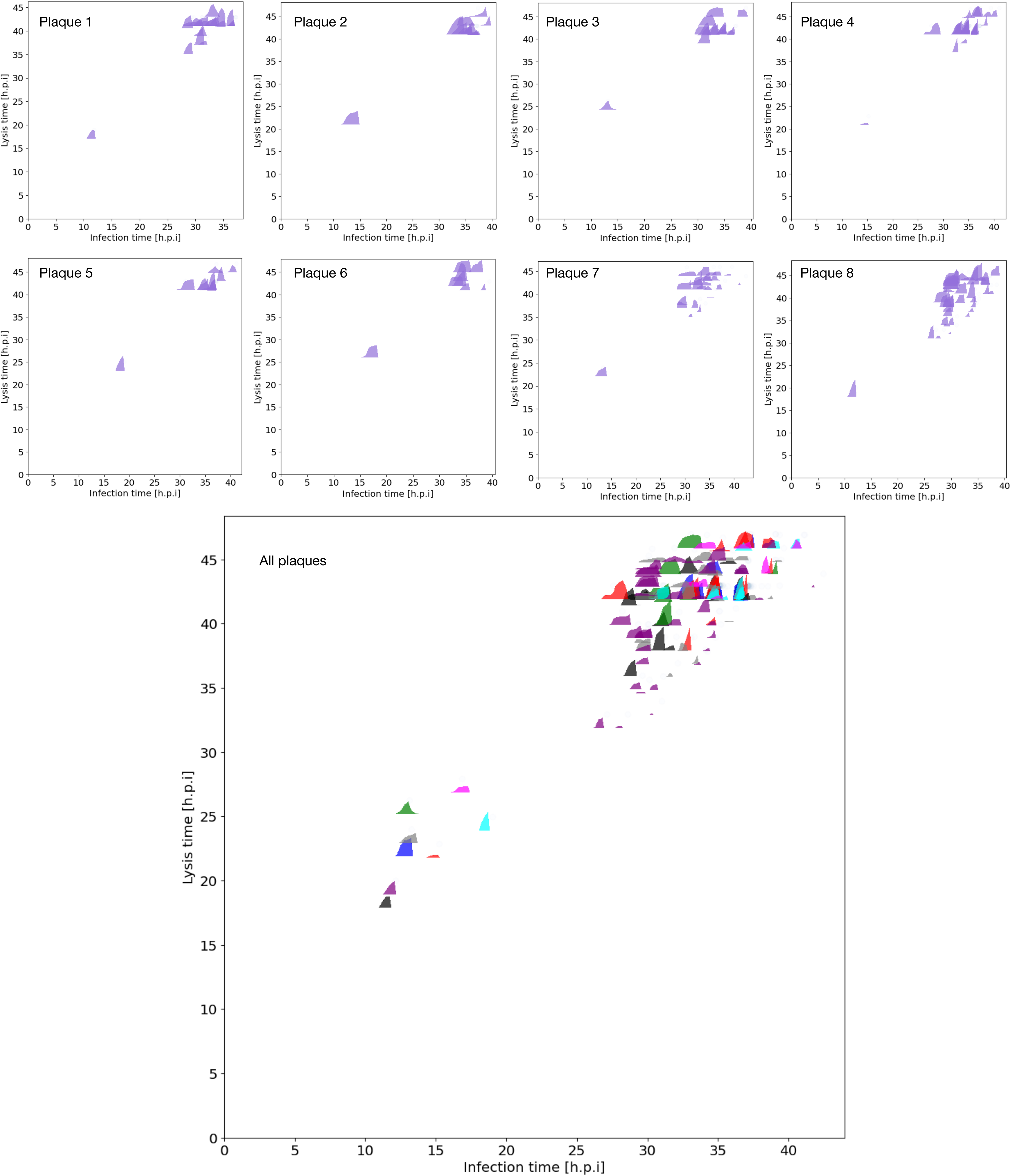
Infection versus lysis time plots for eight plaques are shown, with markers representing individual cells’ intensity profiles. Cells from different plaques with similar clock times are indistinguishable from each other.

Primary infections—defined as the earliest expressing cell in each plaque—appear as early-time outliers, whereas secondary infections form broader clusters whose positions vary across plaques. Across the dataset, the earliest detectable primary infection occurred before 5 hours, and the latest detectable secondary infection continued beyond 40 hours. Given the shared ancestry of secondary infections within a plaque, one might anticipate plaque-specific clusters in infection–lysis timing, but none were initially apparent, underscoring the need for advanced analytical approaches to uncover potential latent relationships.

### Dimensionality Reduction

Linear approaches provided a first view of structure in the dataset. Principal component analysis (PCA) on the standardized feature set revealed that the first three principal components (PCs) accounted for approximately 89.8 percent of total variance (**Fig. 5a**). PC1 was dominated by area under the curve, rise time, infection duration, and GFP max; PC2 captured variation in infection time, rise rate, lysis time, and GFP max; PC3 incorporated additional contributions from lysis time, and rise rate. When projected onto PC2 and PC3 (**Fig. 5b**), primary infected cells formed a sparse cluster of outliers, reflecting their earliest onset times, whereas secondary infected cells clustered more densely together. No clear plaque-wise clustering was observed. The loadings plot further indicated correlations between lysis time and infection time, between area under the curve, GFP max, and rise rate, and between infection duration and rise time. Note that the separation between primary and secondary infected cells aligned with the vectors for infection and lysis times, reflecting the underlying cause of the clusters. Kernel PCA with a sigmoid kernel (γ = 1/n≅0.14) produced similar separations of primary and secondary cells (**Fig. 5c**), with grouping again driven primarily by clock-time attributes rather than plaque identity.

**Figure 5.**
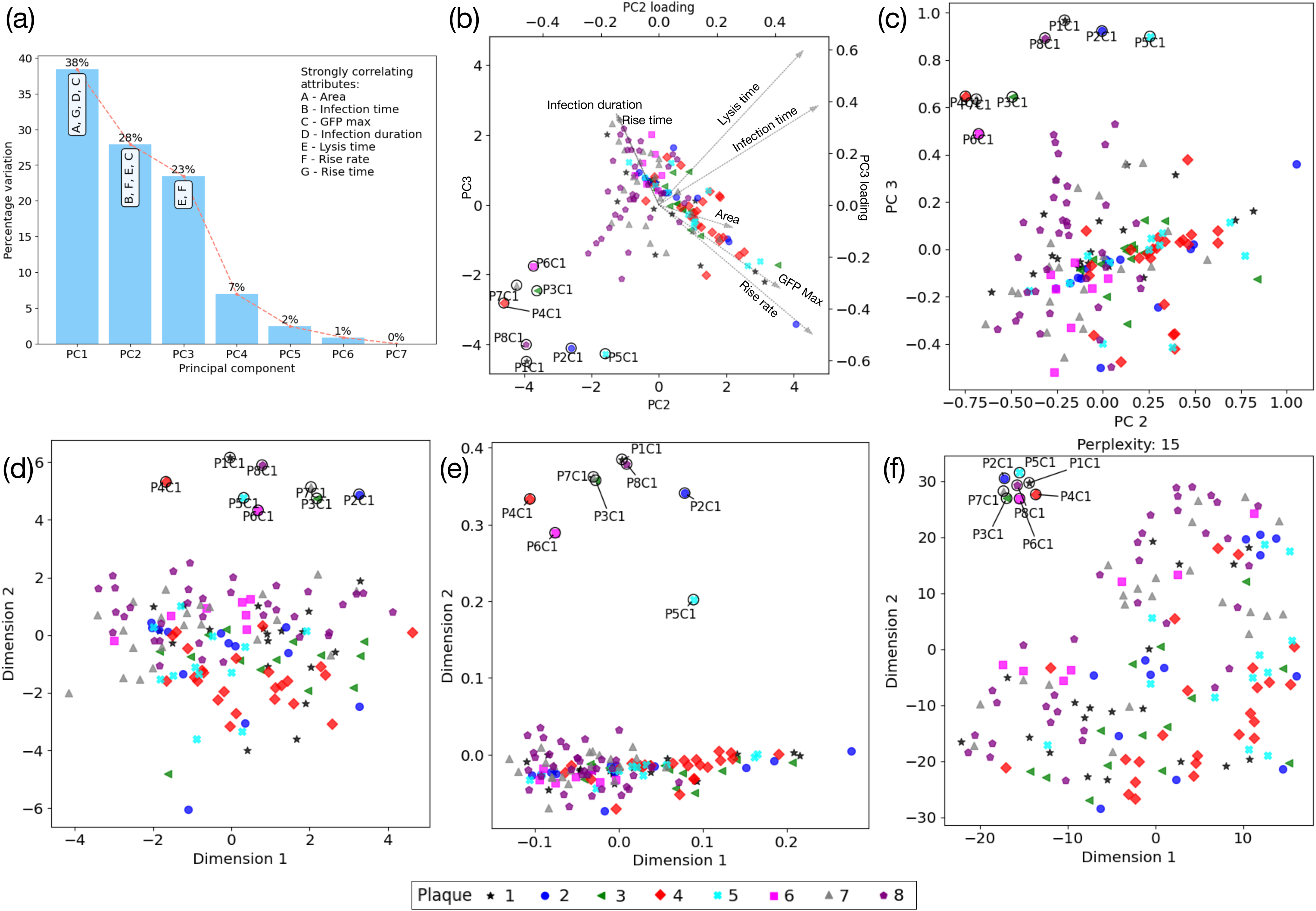
Analysis of experimental data from eight plaques, with 154 cells using dimensionality reduction techniques. In these results, “clusters” refer to groups that appear physically separated in the plots. (a) Eigen values (percentage variance explained) of each principal component (PC). Groups of strongly correlated attributes that are relevant to PCs 1-3 are annotated on their corresponding bars. (b) PCA results: Primary infected cells (annotated as PmC1, where m denotes the plaque number) are the sparse outliers, while secondary infected cells form a dense cluster. Separation is due to different clock-time attributes. Cells within plaques cannot be distinguished as clock-time attributes are relatively close. (c) Kernel PCA (sigmoid, gamma=0.14) produces similar results as PCA: primary cells form a sparse cluster. Secondary cells form a dense cluster with no plaque-wise separation. (d) Metric MDS (2-component) is able to separate primary cells from secondary cells, but unable to capture plaque-to-plaque variations of cells. (e) LLE (2 components, 8 neighbors) of data from eight plaques: primary cells (sparse) are separated from secondary cells (dense cluster). However, it fails to capture plaque-to-plaque variations of cells. (f) t-SNE of data from eight plaques with perplexity=15 separates primary cells (sparse outliers) from secondary cells (dense cluster). However, plaque-wise variations are not captured. Note: The colors and markers assigned to plaques serve the purpose of facilitating cell identification and should not be interpreted as indicative of clustering outcomes. Any cluster labels generated by the method are provided separately.

To examine structure using a geometry-preserving approach, metric multidimensional scaling (MDS) was applied to Euclidean distances between cells in the feature space (**Fig. 5d**). As with PCA and kernel PCA, MDS positioned primary infected cells at the periphery as outliers, but secondary infections largely overlapped in the embedding without forming plaque-specific clusters. This result reinforced the conclusion that similarity in overall timing dominates the variance in the dataset, while plaque-level ancestry contributes little detectable signal at this scale.

Non-linear dimensionality reduction offered complementary perspectives. Locally linear embedding (LLE) with eight nearest neighbors isolated all primary infected cells as outliers (**Fig. 5e**) but collapsed nearly all secondary infected cells into a single dense cluster, failing to resolve plaque-wise distinctions. For t-distributed stochastic neighbor embedding (t-SNE) with perplexity 15 separated secondary infected cells into a distinct cluster (**Fig. 5f**) but again failed to group secondary cells by plaque. These results confirmed that dimensionality reduction can separate primary from secondary infections. However, when infection onset times were similar, the methods failed to reveal plaque-specific signatures.

### Clustering

When density-based clustering (DBSCAN) was applied directly to the seven standardized features using parameters Eps = 1.5 and MinPts = 8, two clusters were identified—labeled −1 and 0 (Fig. 6a). It should be noted that Fig. 6a plots infection time versus lysis time, and the apparent physical separation in this plot does not reflect the clustering outcome. All primary infected cells (annotated in the figure) were classified as outliers (label −1). However, 17 secondary infected cells (appearing in the upper-right quadrant in Fig. 6a) were also included in this outlier category. No plaque-wise separation was evident, as the remaining secondary infected cells from all plaques were mixed together in cluster 0.

**Figure 6.**
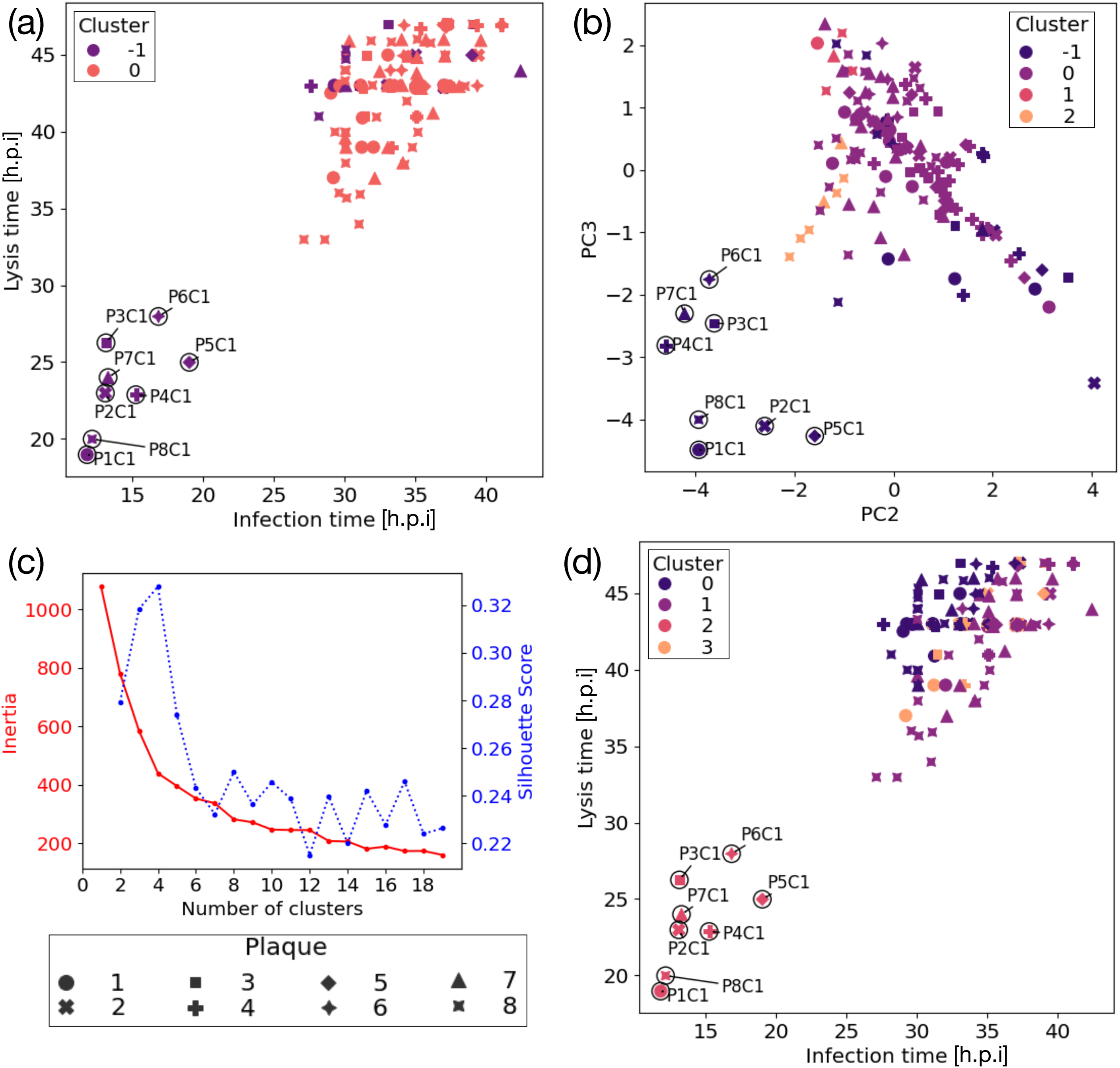
Analysis of experimental data using clustering techniques. Clusters identified by different techniques are conceptually different from those in the previous visualizations, where “clusters” simply referred to groups that appeared physically separated in plots. (a) DBSCAN identifies two clusters. All primary infected-cells are clustered under the outlier cluster (-1). However, 17 secondary cells, which appear physically separated in the infection time vs lysis time plot are also grouped under that cluster. Plaque-to-plaque variations of cells are not captured by this technique as evidenced by cluster 0 which contain cells from multiple plaques. (b) PCA-DBSCAN identified four clusters. All primary cells are clustered under the outlier cluster (-1). However, nineteen secondary cells were misclassified under the outlier cluster. Clusters 0-2 contained a mix of secondary cells from various plaques. (c) Elbow plot of the sum of the squared Euclidean distances of each data point to its closest centroid for a range of K (cluster) values. The slope changes abruptly at K∼4. The silhouette coefficient plot has a maximum at K=4. (d) K-means clustering results: all primary cells are grouped together under cluster 2. Secondary cells from various plaques are mixed in clusters 0, 1 and 3.

Combining PCA with DBSCAN (PCA-DBSCAN) has been shown to be more effective in clustering and outlier detection (22). In our analysis, PCA-DBSCAN identified four clusters (**Fig. 6b**). All primary infected cells were grouped in the outlier cluster (label −1). However, nineteen secondary infected cells were also misclassified into this outlier category. The remaining clusters (labels 0–2) contained mixed populations of secondary cells from different plaques, with 115, 5, and 7 cells in clusters 0, 1, and 2, respectively.

K-means clustering, with K determined by the elbow method and silhouette analysis (**Fig. 6c**), achieved its best separation at K = 4 (silhouette coefficient ≈ 0.33). At this setting (**Fig. 6d**), all eight primary infected cells grouped together correctly (cluster 2), but secondary infected cells from different plaques were mixed into three clusters—0, 1 and 3 with 46, 71, and 29 cells, respectively. These observations indicate that clustering performance is largely dominated by clock-time attributes, with limited sensitivity to plaque-specific differences.

### Sequence-Based Similarity

Sequence comparison methods were applied directly to standardized intensity profiles rather than to extracted features. Dynamic time warping (DTW) computed pairwise distances between profiles, producing a heat map of similarity (**Fig. 7a**) and a dendrogram (**Fig. 7b**). Most profiles appeared highly similar (dark regions in the heat map and short purple edges in the dendrogram), with some light-colored outliers (tall edges in the dendrogram), and no discernible grouping by plaque or infection round. Metric multidimensional scaling (MDS) on the DTW distance matrix (**Fig. 7c**) placed about two-thirds of total cells within 0.2 units from the origin, again indicating high overall similarity, with outliers scattered at greater distances.

**Figure 7.**
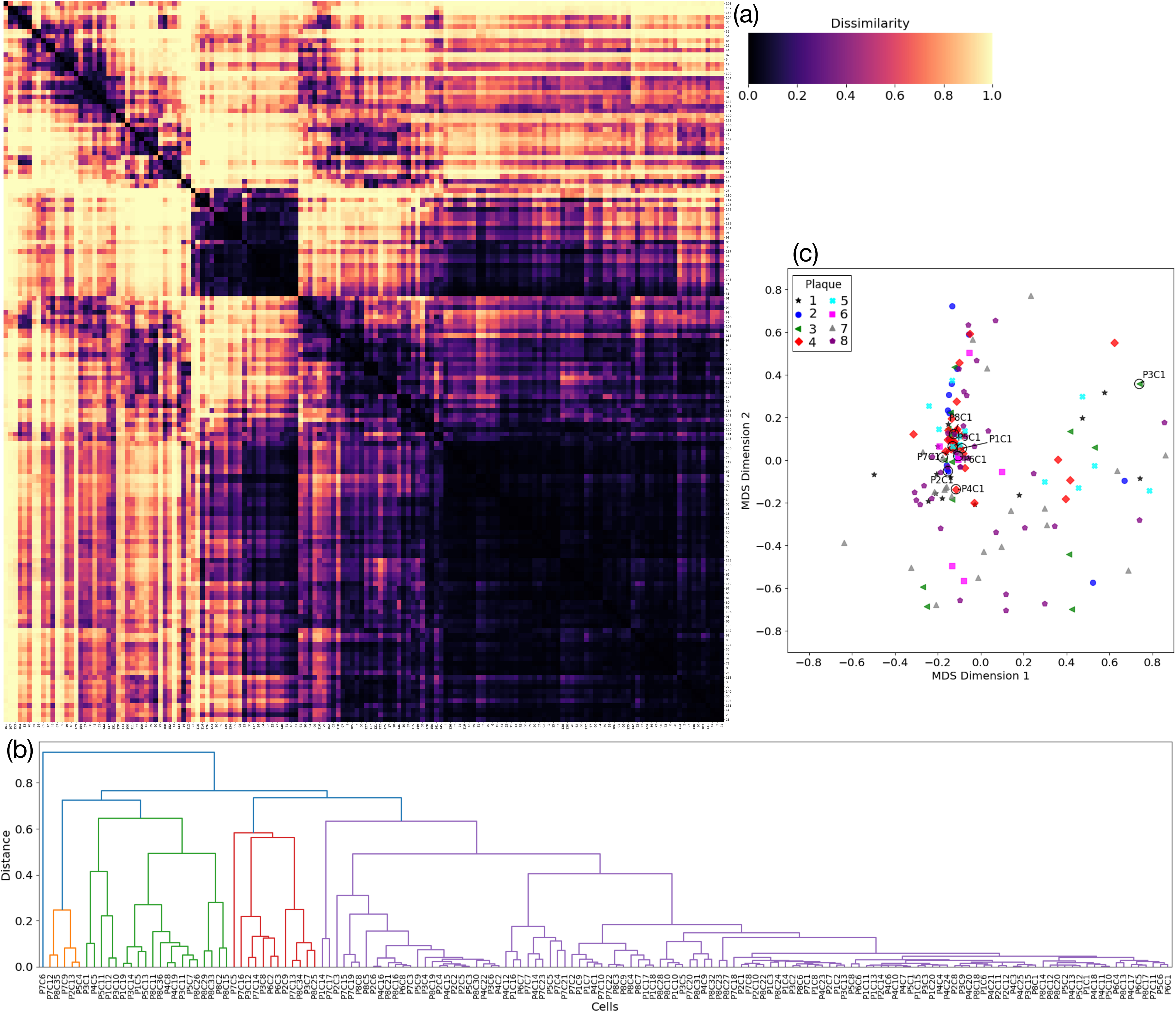
Analysis results of experimental data using DTW: (a) Heat map showing the proximity matrix based on intensity profiles of 154 cells from eight plaques. The color bar indicates the DTW distance which is a measure of dissimilarity between a pair of cells’ intensity profiles. Row and column labels are cell numbers: Plaque 1 (1–20), Plaque 2 (21–34), Plaque 3 (35–49), Plaque 4 (50–74), Plaque 5 (75–87), Plaque 6 (88–95), Plaque 7 (96–118), and Plaque 8 (119–154). Dark on the color bar indicates that the cells’ intensity profiles are similar (i.e., cell-to-cell DTW distance is zero), and light on the color bar indicates that the cells’ profiles are dissimilar (i.e., cell-to-cell DTW distance is one.) Rows and columns have been clustered based on similarities to draw the dendrograms. (b) Dendrogram of all the cells from eight plaques also show that there are no plaque-wise distinguishable clusters. (c) Metric MDS plot in two dimensions using DTW distance matrix for all eight plaques. Cells that are closer on the plot have more similar intensity profiles than those that are further apart. There are no discernible clusters based on plaques or rounds of infection.

The RMSE method, which aligns profiles by shifting in fixed time increments without warping, yielded even greater apparent similarity between cells (**Fig**. **8a**). Metric MDS on the RMSE distance matrix (**Fig. 8b**) and the associated dendrogram (**Fig. 8c**) revealed no clear plaque-wise or round-wise clusters. The lack of separation in both DTW and RMSE analyses suggests that intensity profile shapes are broadly conserved across plaques and rounds, with variability in timing rather than in overall profile form.

**Figure 8.**
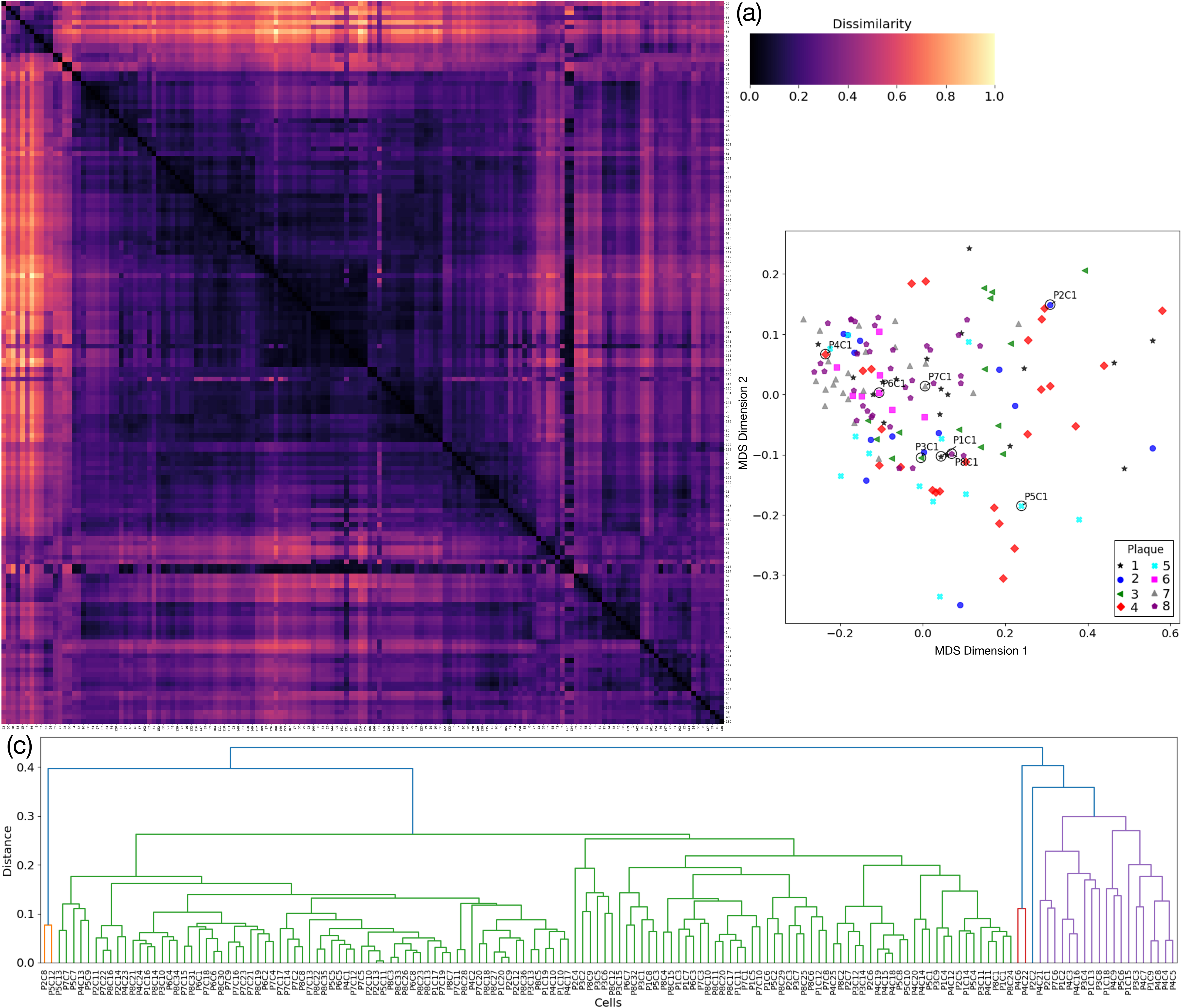
Analysis results of experimental data using the RMSE method: (a) Heat map showing the proximity matrix based on intensity profiles of 154 cells from eight plaques. The color bar indicates the measure of dissimilarity between a pair of cells’ intensity profiles. Row and column labels are cell numbers: Plaque 1 (1–20), Plaque 2 (21–34), Plaque 3 (35–49), Plaque 4 (50–74), Plaque 5 (75–87), Plaque 6 (88–95), Plaque 7 (96–118), and Plaque 8 (119–154). Dark on the color bar indicates that the cells’ intensity profiles are similar, and light on the color bar indicates that the cells’ profiles are dissimilar. Rows and columns have been clustered based on similarities. (b) Two-component metric MDS using the dissimilarity matrix for all eight plaques. Cells that are closer on the plot have more similar intensity profiles than those that are further apart. There are no discernible clusters based on plaques or rounds of infection. (c) Dendrogram of all the cells from eight plaques also show that there are no plaque-wise distinguishable clusters.

### Integrated Comparison

Across all approaches, separation between round-1 (primary) and round-2 (secondary) infections was consistently observed in methods sensitive to timing attributes, such as PCA, kernel PCA, MDS, LLE, t-SNE, and K-means clustering. However, no method achieved robust plaque-wise clustering, reflecting the dominance of clock-time parameters and the minimal intrinsic differences in profile shape across plaques with similar timing. Sequence-based methods (DTW, RMSE) confirmed that the overall form of intensity profiles is conserved, with variability arising chiefly from temporal shifts.

These findings highlight the central role of temporal variables in shaping the dataset and indicate that, under our experimental conditions, plaque identity leaves little imprint on single-cell expression profiles beyond the primary–secondary distinction. While all infections within a plaque trace back to a single initiating cell, the absence of plaque-level clustering could have been anticipated. Even for cells drawn from the same culture well, infections initiated by individual virions display striking heterogeneity, with yields varying over two orders of magnitude [7]. Further, parameters that describe facets of virus progeny release from individual cells also exhibit broad distributions with latent times, rise times, and rise rates spanning two-, five-and over 100-fold, respectively [13]. These parameters, which govern virus output downstream of gene expression, underscore the intrinsic noise and complexity that obscure plaque-specific signatures.

## CONCLUSIONS

The absence of plaque-specific clustering, despite shared ancestry, highlights how quickly infections diversify at the single-cell level. Such heterogeneity in viral gene expression and output may contribute to viral adaptability and unpredictable infection profiles. These insights advance our understanding of how infection spreads from a single cell to a population and suggest new avenues for probing viral fitness and therapeutic response.

## ACKNOWLEDGEMENTS

We thank Yury Bochkov for providing the reporter rhinovirus used in this study. We are grateful for support from the National Science Foundation (DMS-2151959, MCB-2029281 and CBET-2030750), the National Institutes of Health (OT2OD030524, R01DK133605); the Wisconsin Institute for Discovery, Chemical and Biological Engineering, and the Office of the Vice-Chancellor for Research— at University of Wisconsin-Madison.

## Data and code availability

The data that support the findings of this study are available from the corresponding author upon reasonable request. Code from this study is available from github.com/rramachandr9/virus-gene-expression-analysis

## Author contributions

RR conducted data cleaning, analyses, created the figures and wrote the manuscript; HS conducted the wet-lab experiments and wrote the associated section of the methods; JY supervised the research and edited the manuscript.

