## Supporting Information for "From spark to wildfire: illuminating single-cell origins of virus infection spread and diversification"

**Supporting Information 1**

Ramachandran et al.

### 1. Review of Analysis Techniques

The time series profiles of fluorescent protein intensity from individual infected cells, hereafter referred to as intensity profiles, provide a quantitative readout of the infected cell behavior. Of particular interest are the heterogeneity of profiles within or between plaques. Quantifying the similarity or differences between profiles for different cells may provide insight into changes in viral gene expression from generation to generation. Several techniques are well established to compare/quantify similarity between time series data [2]. Some are based on calculation of some inter-sample distance measure (e.g., Euclidean distance). For others, such as principal component analysis (PCA), one needs to extract meaningful attributes from the time series for each sample, and use the extracted data to investigate similarity between samples. Additionally, manifold learning and clustering methods can be used to investigate underlying patterns and structures in such multidimensional data. Prior studies have used various unsupervised learning techniques to analyze and compare single-cell data [37, 34, 14, 30, 11]. However, to our knowledge, no studies have used such techniques to compare virus expression patterns during plaque growth. Here, we briefly review established unsupervised techniques to compare the intensity profiles. These include clustering, dimensionality reduction, and sequence comparisons.

#### 1.1. Dimensionality Reduction Techniques

**Principal component analysis (PCA) and Kernel PCA:** PCA is a commonly used tool to transform the dimensionality of data while retaining most of the information [21]. PCA is useful when many independent variables exhibit multicollinearity to some degree. Transforming the dimensionality helps to identify patterns in the high-dimensional data that otherwise may be difficult to visualize in 3D space. PCA orders Principal Components (PCs) from the most to the least useful to explain the variation in the data. Similar to HC, PCA cannot be used directly on time series data. Instead, for each sample, values of four or more variables are required (because three or fewer variables can be visualized in 3D space) [10]. Since PCA transformations are linear, it is used if the data are linearly separable. However, if the decision boundaries that separate the data are non-linear, kernel PCA is used. Kernel PCA uses nonlinear kernels to project data into higher dimensional space where decision boundaries can be drawn [26]. A limitation of PCA lies in its inability to differentiate between variance caused by measurement noise and variance arising from underlying signal variations [3].

**Multidimensional scaling (MDS):** MDS is a family of dimension-reduction techniques used to quantify and visualize the similarity or dissimilarity between groups of items such as sets of features or sequences. MDS embeds high-dimensional data into a low-dimensional representation that best preserves pairwise distances between data points, thereby modeling similarity or dissimilarity between data points as distances in a geometric space. MDS requires a square distance matrix; each element of the distance matrix is the distance measure (e.g., Euclidean) between a pair of items. Optimization in MDS involves minimization of the stress, where stress is a measure of the discrepancy between the original dissimilarities or similarities and the distances between points in the low-dimensional space [29, 12].

**Isometric mapping:** Isometric mapping is an extension of the classical MDS, and is designed for nonlinear dimensionality reduction. It creates low-dimensional embedding of high-dimensional data using geodesic distances (length of the shortest path between two points that can be followed while remaining on a surface) to calculate a distance matrix. Isometric mapping constructs a neighborhood graph where the edges represent these approximate geodesic distances, and then uses classical MDS to map the data into a lower-dimensional space while preserving these distances as faithfully as possible [28].

**Locally-linear embedding (LLE):** LLE is a nonlinear dimensionality reduction technique that maps high dimensional data into a low dimensional embedding while preserving the local structure of the data [24, 9]. First, each data point is assigned its k nearest neighbors, where k is a hyperparameter. Next, each data point is approximated as a linear combination of these k nearest neighbors. The goal is then to discover a lower-dimensional representation of the dataset that optimally maintains these local relationships. This is done by minimizing the reconstruction error, which quantifies how accurately each data point can be reconstructed using its neighboring points. By relying on this approach, which reconstructs points based on their closest neighbors, the method effectively captures highly nonlinear relationships that characterize the dataset as a whole. However, LLE performs poorly in case of noisy data and outliers [33].

**t-distributed stochastic neighbor embedding (t-SNE):** t-SNE [31] is a nonlinear dimensionality reduction technique for embedding high-dimensional data into a low-dimensional space. It can capture the structure of the underlying data at different scales — local structure as well as global clusters. This is achieved by a hyper parameter called perplexity which determines the width of the Gaussian kernel used to measure similarities between points. As a rule of thumb perplexity is chosen to be smaller than the sample size of the data; perplexity has typical values between 5 and 50. Smaller values of perplexity results in the algorithm focusing on immediate neighbors, and thus emphasizing on local structure. This is useful to capture detailed and fine-grained local relationships. High perplexity values result in the algorithm taking into account a larger number of neighbors, thus capturing more global structure and relationships. That is useful for understanding broader patterns in the data [14].

#### 1.2. Clustering Techniques

**K-means clustering:** K-means clustering is an unsupervised multivariate statistical technique used for partitioning a dataset into K distinct, non-overlapping clusters based on their underlying structure [13, 17, 4]. The number of clusters (K) and the starting centroids are provided as input parameters. This is followed by an iterative process of assigning the data points to each cluster based on their Euclidean squared distance to the corresponding centroid, and recalculating the centroids, until the centroids can no longer be adjusted.

**Hierarchical clustering (HC):** HC is an unsupervised technique that is used to build a hierarchy of clusters of samples based on some inter-sample distance measure (e.g., Euclidean distance, Manhattan distance). HC can be agglomerative (bottom-up approach), or divisive (top-down). In agglomerative HC, we define each data point as a cluster and combine existing clusters at each subsequent step based on linkage criteria. Typical approaches are single, complete, average and centroid linkages, and Ward variance minimization algorithm [27, 20]. Tree-like diagrams called dendrograms are used to represent the clusters graphically. The height of the branches in the dendrogram indicates the distance or dissimilarity at which clusters were merged (for agglomerative HC) or split (for divisive HC). In a dendrogram, the individual data points (and thus the smallest clusters) are called leaves. Leaves are joined by branches (links), whose lengths indicate the dissimilarity between the clusters being joined. Branches meet at nodes and represent clusters at various levels of hierarchy. If several leaves are connected by short branches, then they would form a cluster of highly similar data points. If long branches appear at the lowest level of the dendrogram, then that would suggest that the data points are dissimilar.

**Density-based clustering techniques:** Density-Based Spatial Clustering of Applications with Noise (DBSCAN) is a density-based clustering algorithm which relies on minimum density level estimation based on a threshold for the number of neighbors (MinPts) within an arbitrary distance measure (Eps). It finds areas meeting this minimum density criterion, distinguishing them from regions with lower densities [6, 25, 16]. Its distinct advantage over methods like k-means is its ability to uncover clusters with arbitrary shapes (e.g., nested clusters in high-dimensional spaces.) [8] Hierarchical DBSCAN (HDBSCAN) [5] is an improved algorithm that addresses the major shortcoming of DBSCAN, namely its inability to detect clusters of different densities and handle noisy data effectively. HDBSCAN requires as input the minimum numbers of samples to form a cluster which makes it more intuitive than DBSCAN. Clusters smaller than that will be identified as noise. Ordering points to identify the clustering structure (OPTICS) is yet another density-based clustering technique that can identify clusters of varying densities [1].

**Affinity propagation:** Affinity propagation is a clustering algorithm that does not require specifying the number of clusters in advance. It identifies a set of representative data points, called exemplars, which serve as the centers of clusters. Initially, every data point is considered a potential exemplar. The algorithm uses a measure of affinity, which is often based on the negative of the Euclidean distance between points, to evaluate the likelihood of each point being an exemplar. Higher affinity indicates a stronger likelihood that a data point will be chosen as an exemplar. The final clusters are formed by grouping data points around these exemplars, with the algorithm iteratively refining the assignment of points to clusters until convergence [7, 22].

**Spectral Clustering:** Spectral clustering differs from other clustering methods by using the eigenvalues and eigenvectors of a similarity matrix derived from the data to perform dimensionality reduction before clustering it in a lower dimensional space. It is able to cluster non-linear data and complex cluster shapes [19].

#### 1.3. Sequence Comparison Methods

**Dynamic time warping (DTW):** DTW is a technique used to compare and quantify the similarity between two time-dependent sequences of different durations or speeds. It does so by changing the alignment of data points. It is commonly used in domains such as speech and gesture recognition, financial analysis, and medicine owing to its robustness [18]. While DTW often finds a global optimum solution, it does not always yield locally sensible matchings. Specifically, DTW may pair two temporal points with entirely dissimilar local structures due to over-stretching and over-compression [36, 35, 15, 23].

**Cross-correlation:** Cross-correlation is a technique used to measure the similarity between two time series as a function of the displacement of one relative to the other. Cross correlation requires paired data [32]. Thus, for sequences of unequal lengths, the shorter sequence needs to be padded, or the longer sequence needs to be truncated.

[9] Benyamin Ghojogh, Ali Ghodsi, Fakhri Karray, and Mark Crowley. Locally linear embedding and its variants: Tutorial and survey, 2020.

[27] Robert R Sokal and Charles D Michener. A statistical method for evaluating systematic relationships. University of Kansas Scientific Bulletin, 38, 1958.

#### Table 1: Summary of Clustering Methods

| Method | Key Principle | Input Requirements | Advantages | Limitations | Best Suited For |
|---|---|---|---|---|---|
| K-means Clustering | Partitions data into K clusters by minimizing intra-cluster variance | Number of clusters (K), initial centroids | Simple, fast, works well for spherical clusters | Sensitive to K, poor with non-globular or noisy data | Data with clear, spherical cluster boundaries |
| Hierarchical Clustering | Builds a tree of clusters using linkage criteria | Distance metric, linkage method | Does not require number of clusters a priori, dendrogram visualization | Computationally expensive, sensitive to noise | Small to medium datasets with hierarchical relationships |
| DBSCAN | Groups points based on density (MinPts within Eps radius) | Eps (distance), MinPts | Detects arbitrary shapes, handles noise | Struggles with varying densities, sensitive to parameter choice | Spatial data, clusters of different shapes |
| HDBSCAN | Extends DBSCAN to handle varying densities via hierarchical clustering | Min cluster size | Better for noisy and variable-density data | May produce many small clusters or noise | Noisy data, biological datasets with unclear boundaries |
| OPTICS | Orders data to identify clustering structure | Eps, MinPts | Handles varying densities, visual hierarchy of clustering | Complexity, harder interpretation | Large datasets with unknown cluster structure |
| Affinity Propagation | Uses message-passing to identify exemplars and form clusters | Preference, damping factor | No need to pre-specify K, robust to initial conditions | High computational cost, sensitive to input preference parameter | Situations where exemplars are natural representatives |
| Spectral Clustering | Uses graph Laplacian of similarity matrix, clusters in reduced eigenspace | Similarity matrix | Effective for non-linear clusters and complex shapes | Sensitive to similarity matrix, computationally intensive | Clustering in non-linear manifolds, image and gene expression data |

#### Table 2: Summary of Dimensionality Reduction Methods

| Method | Key Principle | Input Requirements | Advantages | Limitations | Best Suited For |
|---|---|---|---|---|---|
| PCA | Linear transformation to maximize variance via orthogonal principal components | Numeric data, standardized features | Simple, interpretable, fast | Only captures linear relationships, sensitive to scaling | Preliminary data exploration, noise filtering |
| Kernel PCA | Applies PCA in high-dimensional space via kernel trick | Kernel choice (e.g., RBF, sigmoid) | Captures non-linear patterns | PCs hard to interpret, sensitive to kernel parameters | Complex datasets with non-linear relationships |
| MDS | Embeds high-dim data into low-dim space while preserving pairwise distances | Distance/dissimilarity matrix | Intuitive similarity visualization | Stress optimization sensitive, cannot handle missing distances | Visualization of perceptual/ biological similarities |
| Isomap | Nonlinear extension of MDS using geodesic distances on nearest-neighbor graphs | Neighborhood size (k or Eps) | Preserves global structure in curved manifolds | Sensitive to neighborhood parameter, fails with disconnected manifolds | Nonlinear structure with global patterns |
| LLE | Preserves local linear relationships among neighbors | Number of neighbors (k) | Good at capturing local structure, nonlinear relationships | Poor noise tolerance, hard to tune | Local pattern discovery in noisy biological data |
| t-SNE | Embeds data using pairwise probabilities to maintain local similarities | Perplexity (typically 5–50) | Captures both global and local clusters visually | Not deterministic, poor with large-scale distances | Visualization of high-dimensional single-cell or transcriptomic data |

**Supporting Information 2**

Ramachandran et al.

We analyzed the various techniques discussed in the manuscript using the toy datasets S1, S1-10, S1-20, S1-30, S1-40, S2 and S3.

Toy dataset S1 consists of four plaques, each with ten identical intensity profiles. There is no cell-to-cell variability of profiles within any plaque, whereas there is plaque-to-plaque dissimilarity as the profiles from each plaque have distinct shapes. The round-2 cells' profiles are evenly spaced out in time. S1 is meant to replicate the effect of genetic variability between plaques on the analyses. Toy dataset S2 consists of four plaques, each with ten similar intensity profiles. S2 is similar to S1 but there is cell-to-cell variability of profiles within any plaque as the profiles are similar but of different scales. Also, round-2 cells' profiles are randomly spaced out in time. S2 is meant to replicate the effect of cell-to-cell environmental and genetic variability within and between plaques on the analyses. We introduced Gaussian random noise at various percentages (10%, 20%, 30%, and 40% of peak intensity) to create datasets S1-10, S1-20, S1-30, and S1-40. These datasets allow us to study the impact of noise on our analyses.

Additionally, we generated toy dataset S3, which comprises six plaques with varying cell counts (ranging from 6 to 13). Each plaque has randomly generated intensity profiles, simulating plaques with dissimilar cells. Dataset S3 is designed to test the case where only clock time attributes differentiate between round-1 and round-2 cells. Other similarities observed in the results are by random chance.

As some of these techniques cannot be applied directly to compare time series of unequal lengths, we first extracted several features/attributes from the time series profiles as described in the main text. We scored each method based on whether it was able to identify or separate round-1 cells of each plaque from the rest, and cluster round-2 cells plaque-wise. The results are presented in Tables 1, 2, 3, and 4. DBSCAN, PCA-DBSCAN, kernel PCA, and metric MDS scored the highest in their ability to separate round-1 cells (see Table 1). K-means clustering, spectral clustering, affinity propagation, and t-SNE scored the highest in their ability to separate round-2 cells (see Table 2). Considering noise in the toydata (see Table 3), K-means clustering, t-SNE, metric MDS scored the highest. When considering only the dataset S2 (with cell-to-cell variability of profiles within any plaque), DTW, scored the highest (see Table 4.)

The results of analysis of dataset S3 reveal that clock time attributes have a strong influence on the clustering outcomes for LLE and t-SNE, with round-1 cells distinctly clustered together, separate from round-2 cells. This trend is also observed, though to a slightly lesser extent, in spectral clustering, PCA, kernel PCA, MDS, and isometric mapping, where round-1 cells are generally distinct from round-2 cells. DBSCAN shows only a moderate sensitivity to clock time attributes, as some round-2 cells are grouped with round-1 cells. On the other hand, K-means clustering, HDBSCAN, OPTICS, affinity propagation, and PCA-DBSCAN are only very mildly affected by clock time, as round-1 cells are not fully distinguishable from round-2 cells. Meanwhile, DTW and RMSE remain unaffected by clock time, as it is not considered in their analysis. In the case of S3, the only inherent similarity across the plaques is that round-1 cells share similar clock times. Therefore, the stronger the effect of clock time attributes on the results, the better. However, it is important to note that this focus on clock time could potentially cause the overlooking of clusters that might arise from other underlying factors.

**Table 1:** Scores based on ability to cluster round-1 cells.

| Method | S1 | S1-10 | S1-20 | S1-30 | S1-40 | S2 | Score % |
|---|---|---|---|---|---|---|---|
| DBSCAN | 4 | 4 | 4 | 4 | 4 | 4 | 100.00% |
| PCA-DBSCAN | 4 | 4 | 3 | 4 | 3 | 4 | 91.67% |
| Kernel PCA | 4 | 4 | 2 | 4 | 3 | 3 | 83.33% |
| MDS | 4 | 4 | 4 | 4 | 3 | 1 | 83.33% |
| HDBSCAN | 1 | 2 | 4 | 4 | 2 | 4 | 70.83% |
| K-means | 4 | 3 | 3 | 3 | 2 | 1 | 66.67% |
| PCA | 4 | 4 | 2 | 3 | 2 | 1 | 66.67% |
| t-SNE | 4 | 4 | 4 | 4 | 0 | 0 | 66.67% |
| LLE | 4 | 4 | 4 | 3 | 0 | 0 | 62.50% |
| OPTICS | 1 | 2 | 3 | 4 | 3 | 1 | 58.33% |
| Isometric mapping | 3 | 3 | 3 | 3 | 1 | 1 | 58.33% |
| Affinity propagation | 0 | 0 | 0 | 0 | 3 | 0 | 12.50% |
| Spectral clustering | 0 | 0 | 0 | 0 | 0 | 0 | 0.00% |
| DTW | 0 | 0 | 0 | 0 | 0 | 0 | 0.00% |
| RMSE method | 0 | 0 | 0 | 0 | 0 | 0 | 0.00% |

**Table 2:** Scores based on ability to cluster round-2 cells.

| Method | S1 | S1-10 | S1-20 | S1-30 | S1-40 | S2 | Score % |
|---|---|---|---|---|---|---|---|
| K-means | 36 | 35 | 36 | 36 | 33 | 26 | 93.52% |
| Spectral clustering | 36 | 35 | 36 | 36 | 33 | 26 | 93.52% |
| Affinity propagation | 36 | 35 | 36 | 36 | 28 | 20 | 88.43% |
| t-SNE | 36 | 35 | 36 | 36 | 31 | 17 | 88.43% |
| OPTICS | 36 | 27 | 36 | 34 | 32 | 25 | 87.96% |
| Isometric mapping | 36 | 35 | 35 | 36 | 22 | 20 | 85.19% |
| HDBSCAN | 36 | 35 | 36 | 34 | 24 | 16 | 83.80% |
| MDS | 36 | 35 | 36 | 36 | 27 | 10 | 83.33% |
| DTW | 36 | 36 | 34 | 28 | 6 | 36 | 81.48% |
| LLE | 36 | 34 | 35 | 21 | 28 | 15 | 78.24% |
| PCA-DBSCAN | 36 | 28 | 34 | 25 | 19 | 14 | 72.22% |
| RMSE method | 36 | 36 | 34 | 34 | 0 | 13 | 70.83% |
| DBSCAN | 36 | 20 | 32 | 25 | 25 | 8 | 67.59% |
| Kernel PCA | 36 | 35 | 30 | 18 | 16 | 0 | 62.50% |
| PCA | 33 | 32 | 31 | 33 | 0 | 0 | 59.72% |

**Table 3:** Scores based on noise in toy data.

| Method | S1 | | S1-10 | | S1-20 | | S1-30 | | S1-40 | | Score % |
|---|---|---|---|---|---|---|---|---|---|---|---|
| | round -1 | round -2 | round -1 | round -2 | round -1 | round -2 | round -1 | round -2 | round -1 | round -2 | |
| K-means | 4 | 36 | 3 | 35 | 3 | 36 | 3 | 36 | 2 | 33 | 95.50 % |
| t-SNE | 4 | 36 | 4 | 35 | 4 | 36 | 4 | 36 | 0 | 31 | 95.00 % |
| MDS | 4 | 36 | 4 | 35 | 4 | 36 | 4 | 36 | 3 | 27 | 94.50 % |
| HDBSCAN | 1 | 36 | 2 | 35 | 4 | 36 | 4 | 34 | 2 | 24 | 89.00 % |
| OPTICS | 1 | 36 | 2 | 27 | 3 | 36 | 4 | 34 | 3 | 32 | 89.00 % |
| Isometric mapping | 3 | 36 | 3 | 35 | 3 | 35 | 3 | 36 | 1 | 22 | 88.50 % |

| | | | | | | | | | | | |
|---|---|---|---|---|---|---|---|---|---|---|---|
| Spectral clustering | 0 | 36 | 0 | 35 | 0 | 36 | 0 | 36 | 0 | 33 | 88.00 % |
| Affinity propagation | 0 | 36 | 0 | 35 | 0 | 36 | 0 | 36 | 3 | 28 | 87.00 % |
| LLE | 4 | 36 | 4 | 34 | 4 | 35 | 3 | 21 | 0 | 28 | 84.50 % |
| PCA-DBSCAN | 4 | 36 | 4 | 28 | 3 | 34 | 4 | 25 | 3 | 19 | 80.00 % |
| DBSCAN | 4 | 36 | 4 | 20 | 4 | 32 | 4 | 25 | 4 | 25 | 79.00 % |
| Kernel PCA | 4 | 36 | 4 | 35 | 2 | 30 | 4 | 18 | 3 | 16 | 76.00 % |
| PCA | 4 | 33 | 4 | 32 | 2 | 31 | 3 | 33 | 2 | 0 | 72.00 % |

| | | | | | | | | | | | |
|---|---|---|---|---|---|---|---|---|---|---|---|
| DTW | 0 | 36 | 0 | 36 | 0 | 34 | 0 | 28 | 0 | 6 | 70.00 % |
| RMSE method | 0 | 36 | 0 | 36 | 0 | 34 | 0 | 34 | 0 | 0 | 70.00 % |

**Table 4:** Scores based on ability to cluster cells in S2.

| Method | S2 | | Score % |
|---|---|---|---|
| | round-1 | round-2 | |
| DTW | 0 | 36 | 90.00% |
| K-means | 1 | 26 | 67.50% |
| OPTICS | 1 | 25 | 65.00% |
| Spectral clustering | 0 | 26 | 65.00% |
| Isometric mapping | 1 | 20 | 52.50% |
| HDBSCAN | 4 | 16 | 50.00% |
| Affinity propagation | 0 | 20 | 50.00% |
| PCA-DBSCAN | 4 | 14 | 45.00% |
| t-SNE | 0 | 17 | 42.50% |
| LLE | 0 | 15 | 37.50% |
| RMSE method | 0 | 13 | 32.50% |
| DBSCAN | 4 | 8 | 30.00% |
| MDS | 1 | 10 | 27.50% |
| Kernel PCA | 3 | 0 | 7.50% |
| PCA | 1 | 0 | 2.50% |

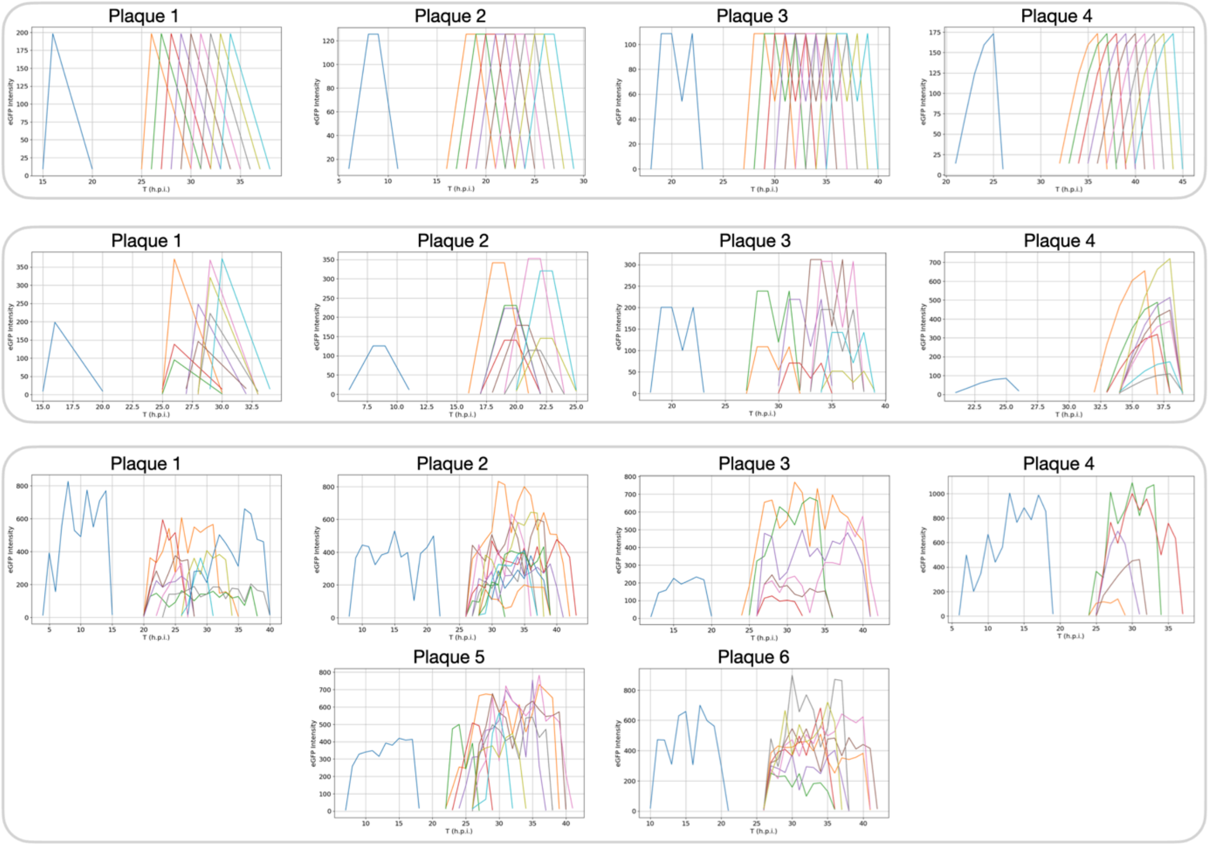


**Figure 1**: Toy datasets S1, S2, and S3. S1 consists of four plaques, each with 10 identical intensity profiles. There is no cell-to-cell variability of profiles within any plaque, whereas there is plaque-to-plaque dissimilarity as the profiles from each plaque have distinct shapes. The round-2 cells' profiles are evenly spaced out in time. S2 consists of four plaques, each with 10 similar intensity profiles. S2 is similar to S1 but there is cell-to-cell variability of profiles within any plaque as the profiles are similar but of different scales. Also, round-2 cells' profiles are randomly spaced out in time. S3 comprises six plaques with varying cell counts (ranging from six to 13) and each plaque has randomly generated intensity profiles.

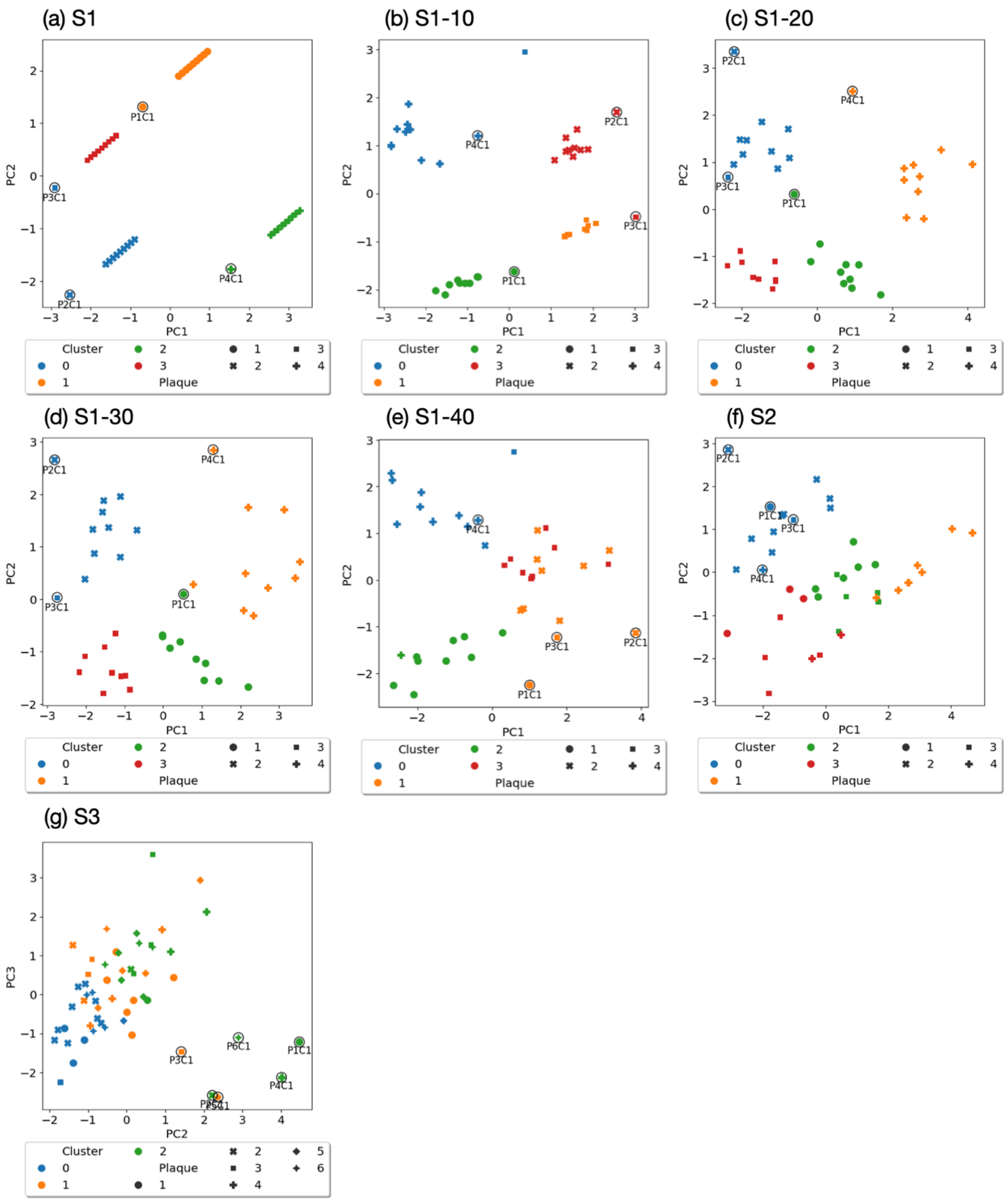


**Figure 2**: K-means analysis of attributes extracted from toy datasets. Data is visualized using principal components 1 and 2.

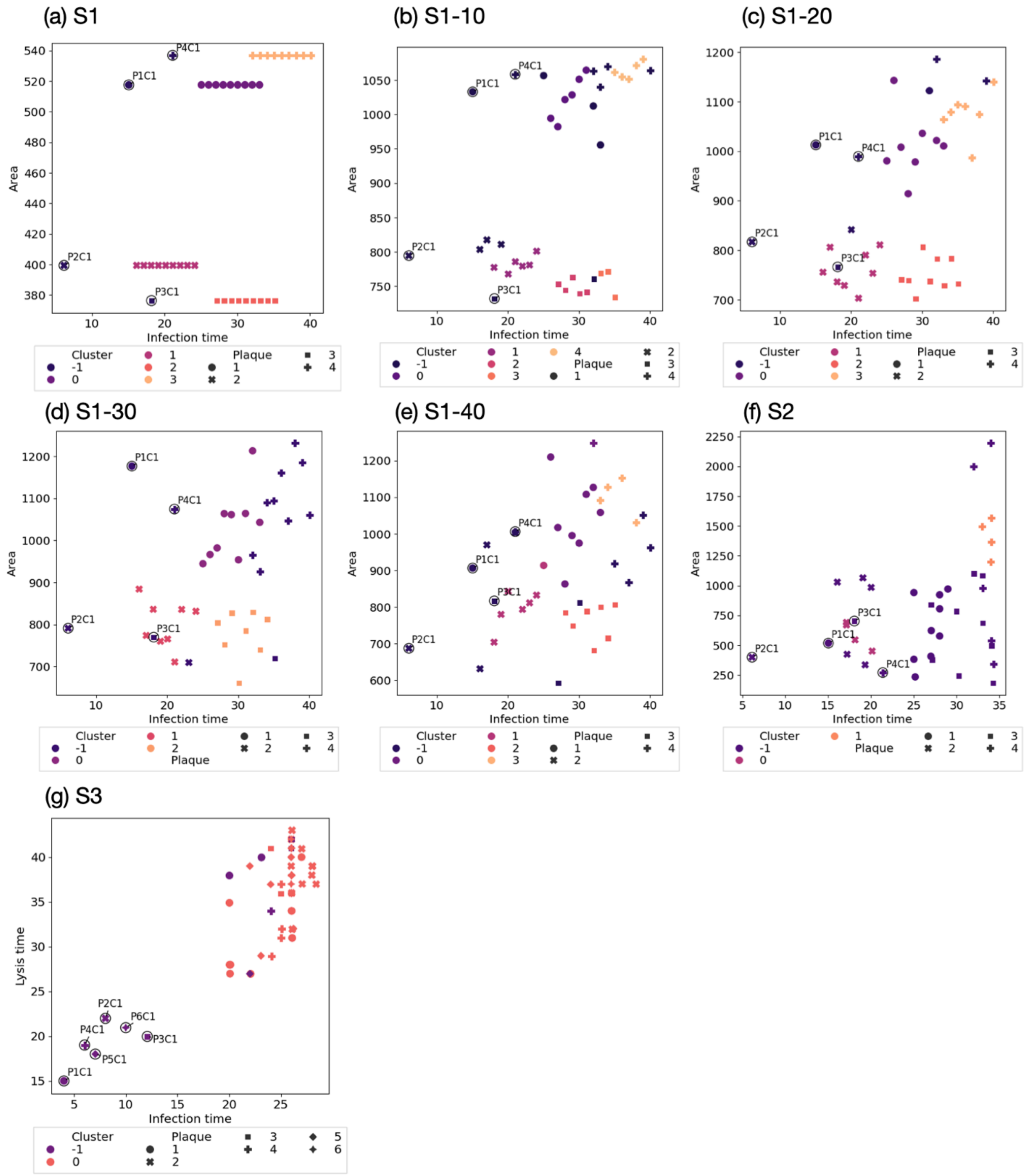


**Figure 3**: DBSCAN analysis of attributes extracted from toy datasets. Cluster -1 are outliers. Units - area [a.u. h], infection time [h.p.i], lysis time [h.p.i].

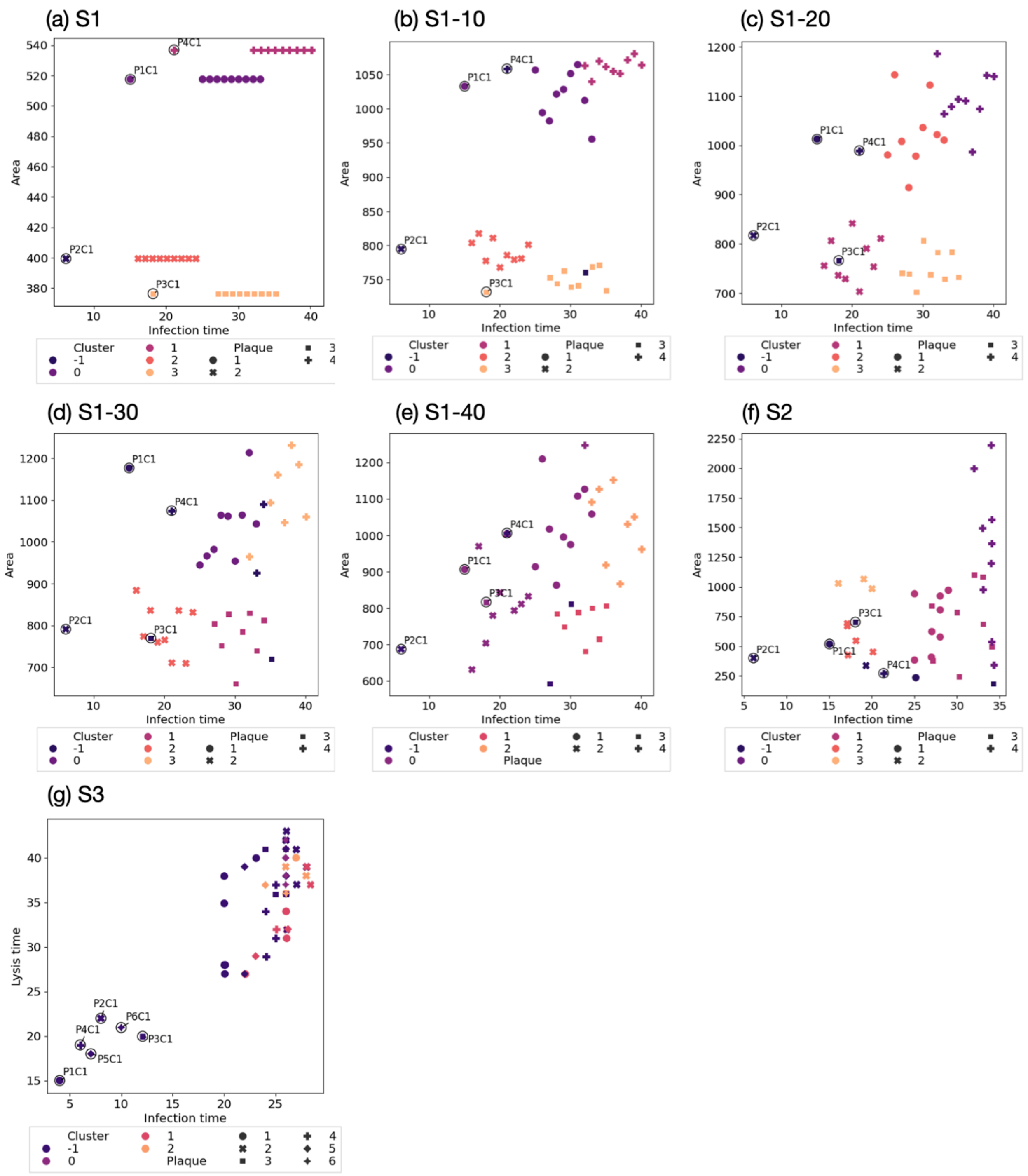


**Figure 4**: HDBSCAN analysis of attributes extracted from toy datasets. Cluster -1 are outliers. Units - area [a.u. h], infection time [h.p.i], lysis time [h.p.i].

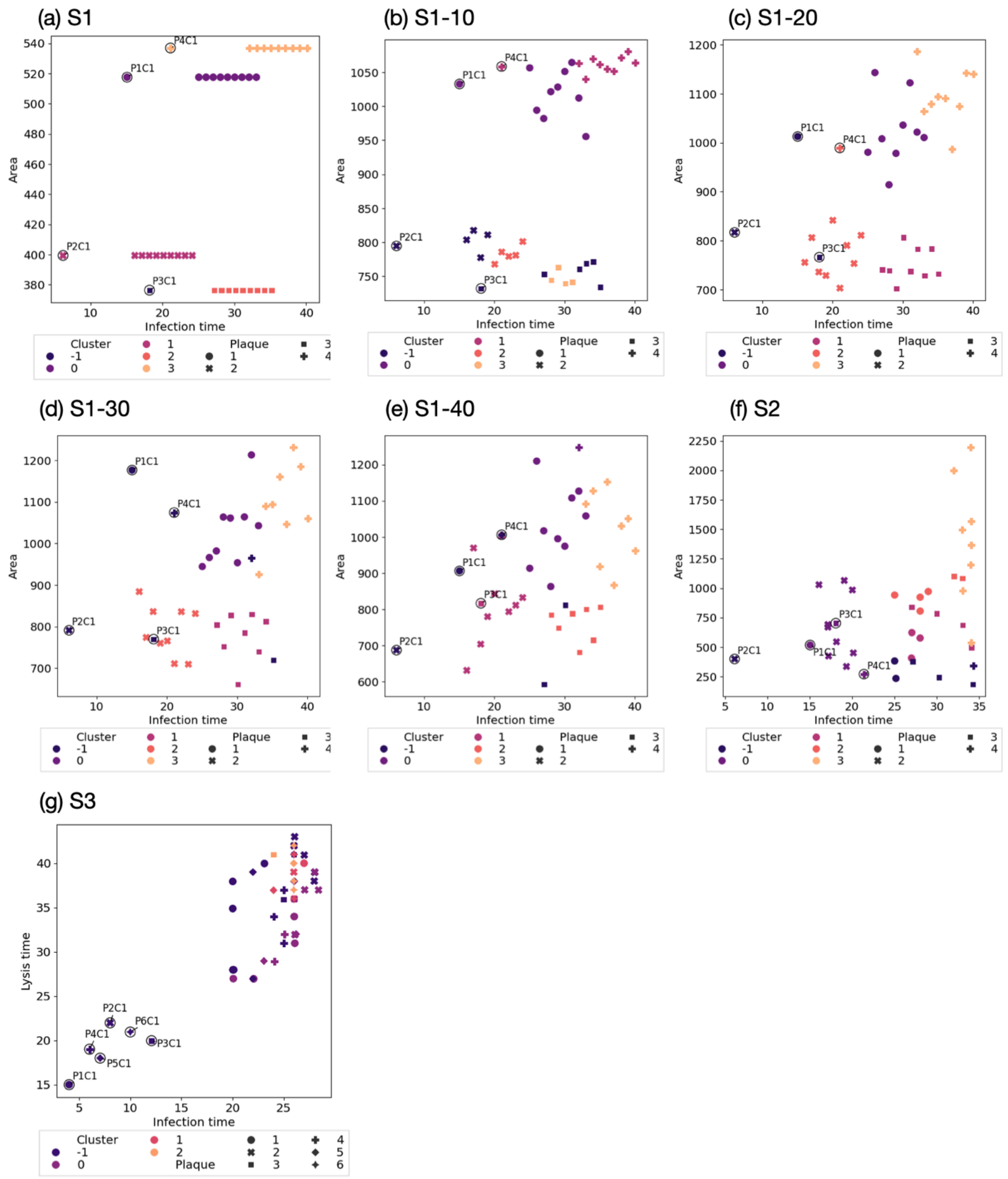


**Figure 5**: OPTICS analysis of attributes extracted from toy datasets. Cluster -1 are outliers. Units - area [a.u. h], infection time [h.p.i], lysis time [h.p.i].

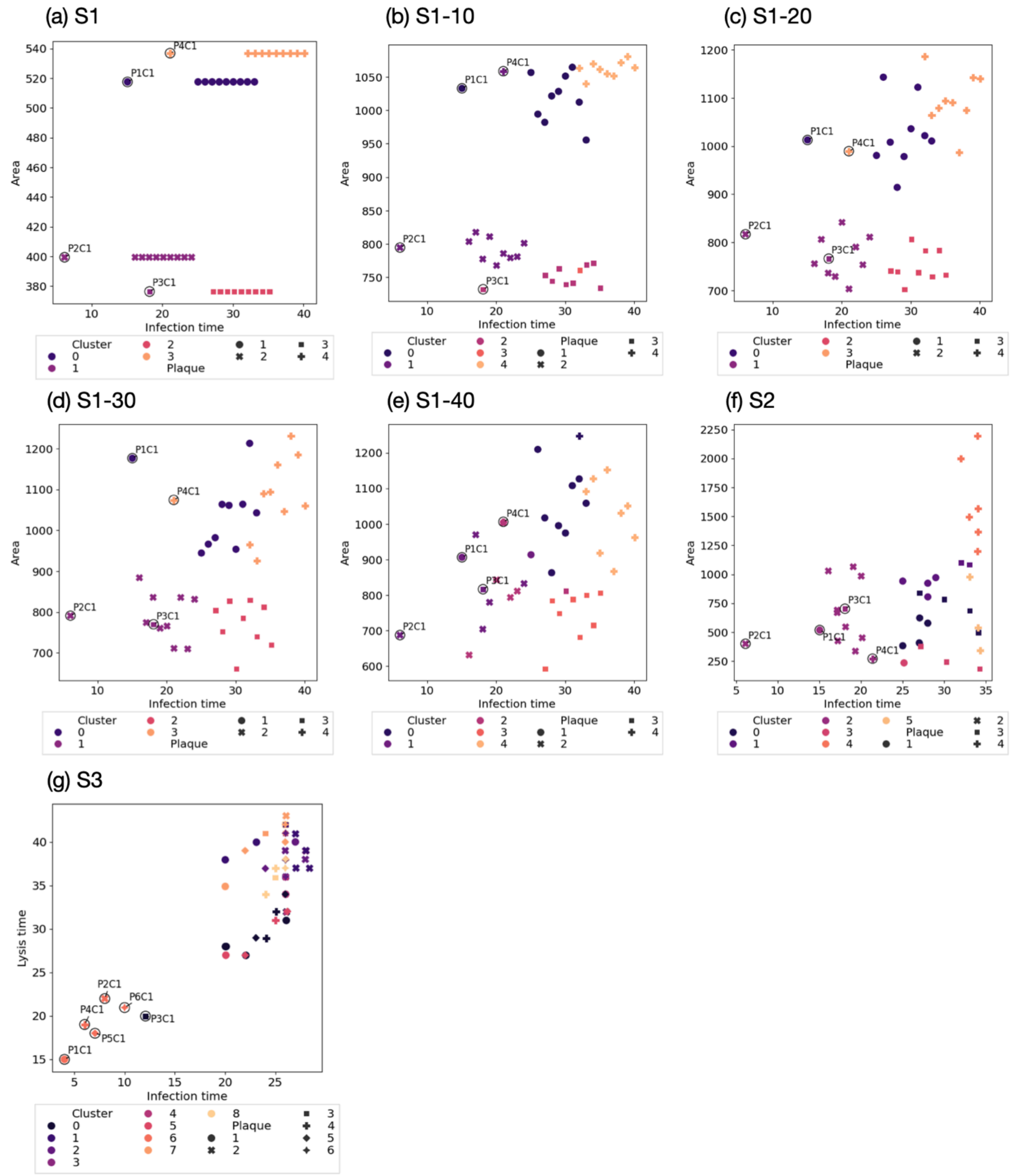


**Figure 6**: Affinity propagation analysis of attributes extracted from toy datasets.

Units - area [a.u. h], infection time [h.p.i], lysis time [h.p.i].

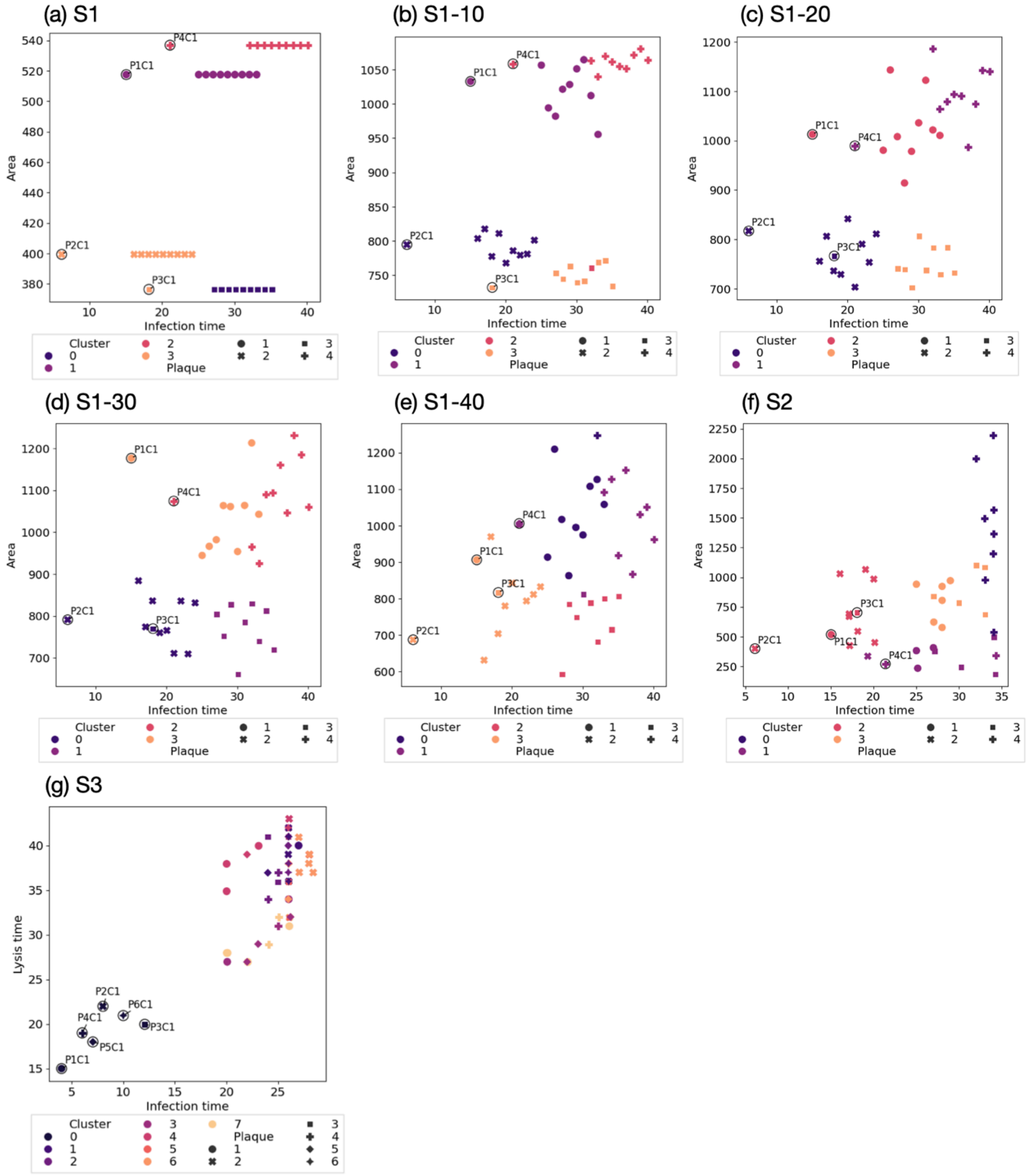


**Figure 7**: Spectral clustering analysis of attributes extracted from toy datasets.

Units - area [a.u. h], infection time [h.p.i], lysis time [h.p.i].

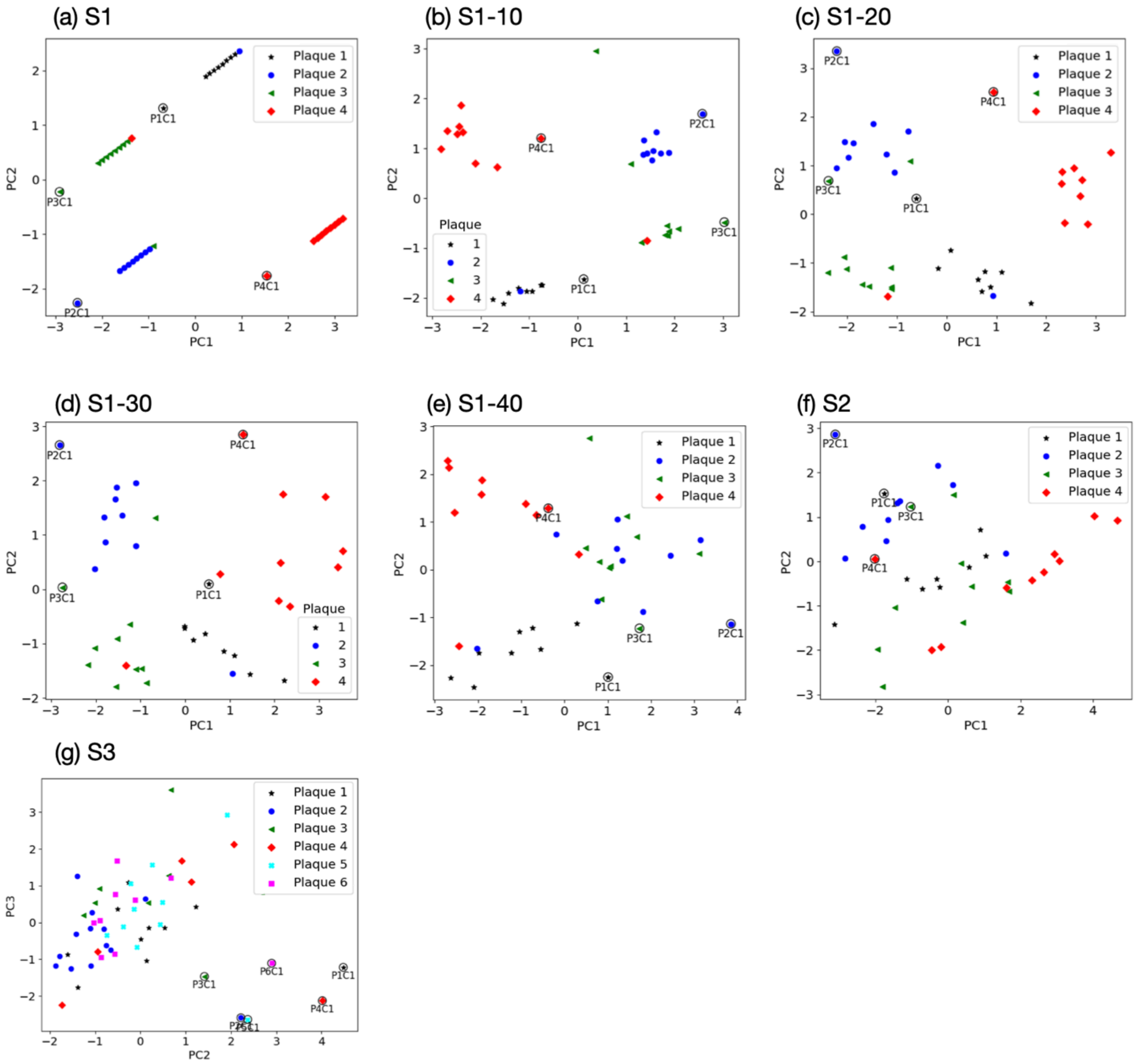


**Figure 8**: Principal component analysis of attributes extracted from toy datasets.

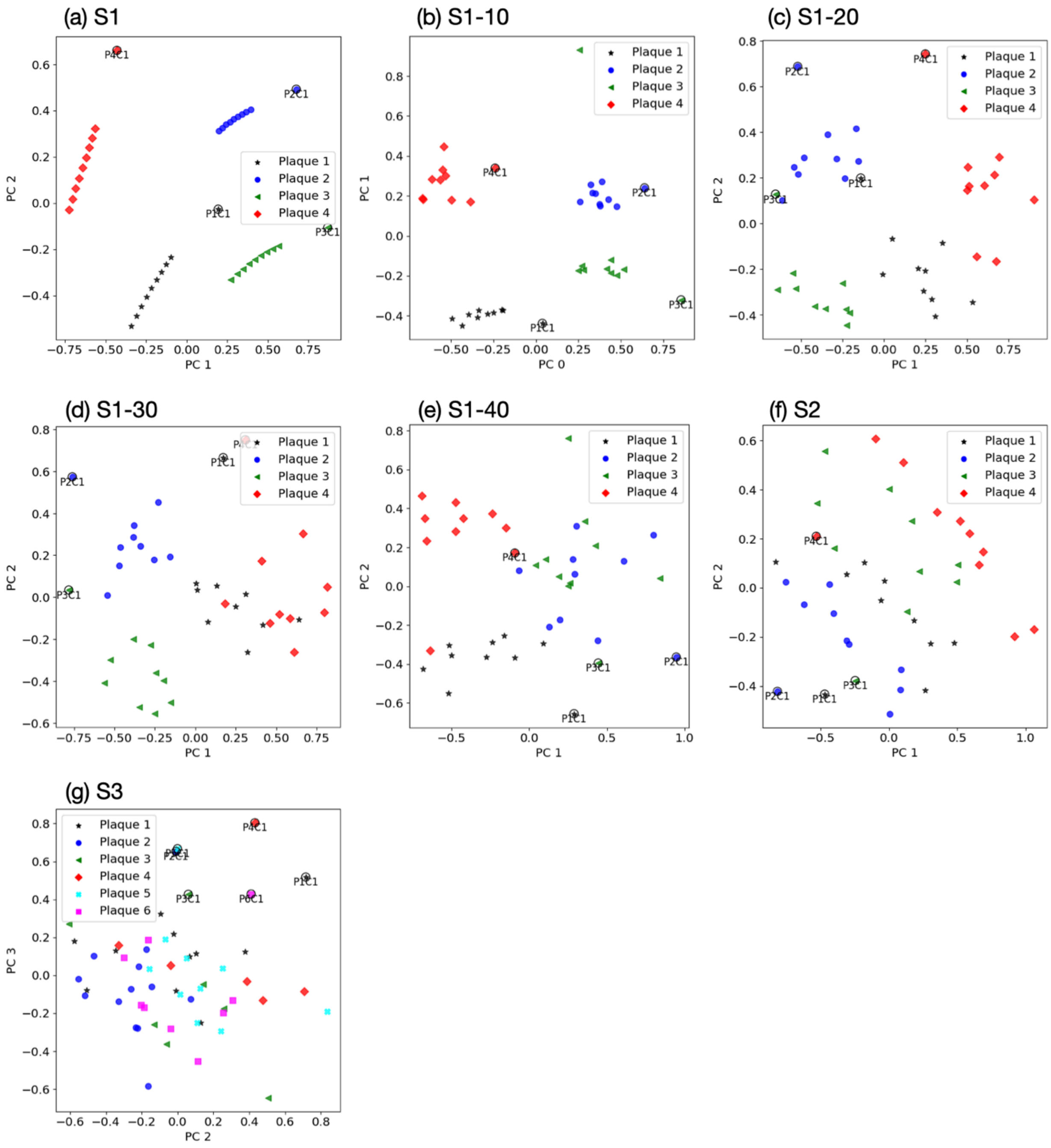


**Figure 9:** Kernal PCA of attributes extracted from toy datasets.

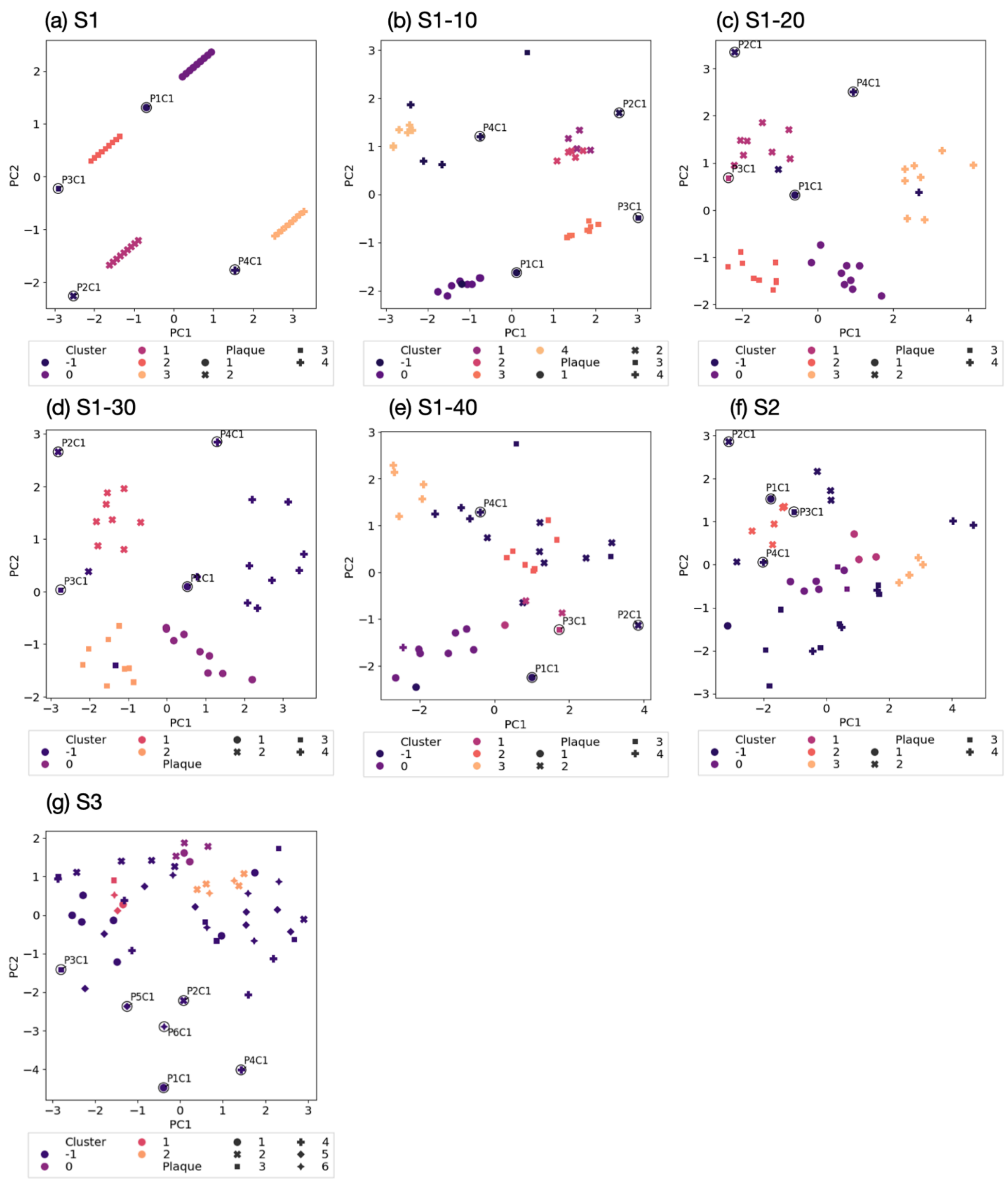


**Figure 10:** Analysis of toy datasets' attributes using PCA-DBSCAN. Cluster -1 are outliers

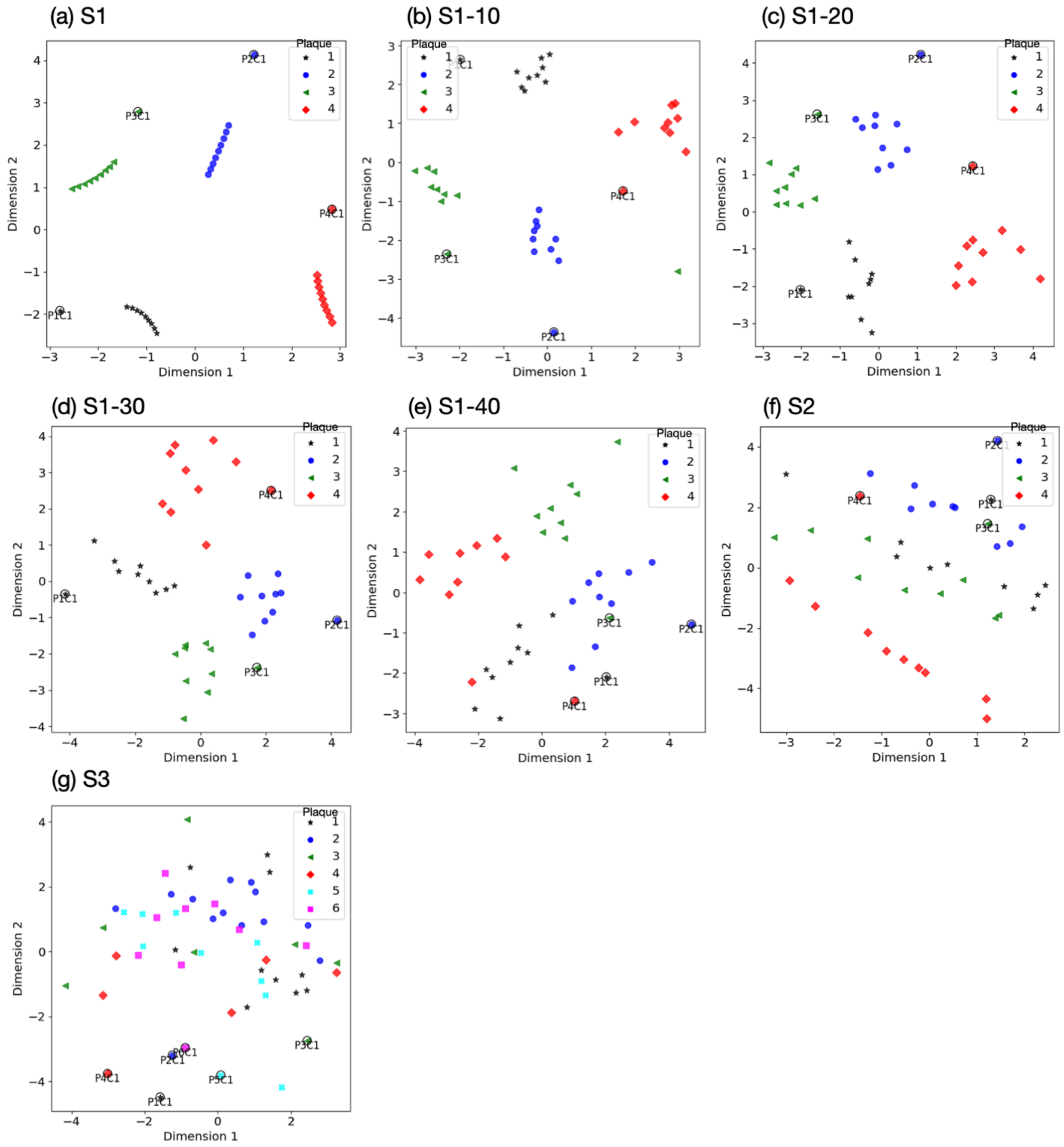


**Figure 11:** MDS analysis of toy datasets' attributes

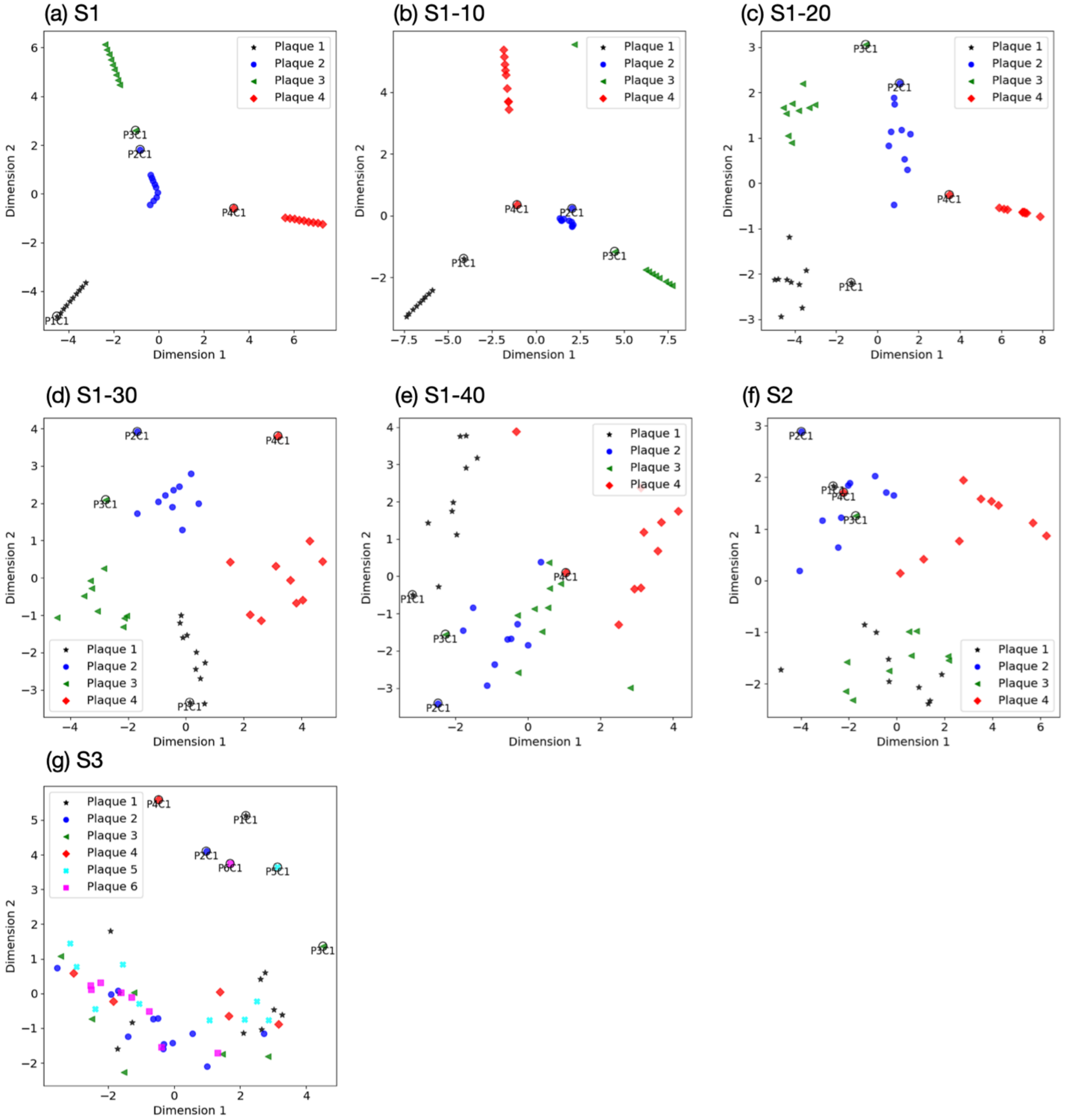


**Figure 12:** Analysis of attributes extracted from toy datasets using isomap

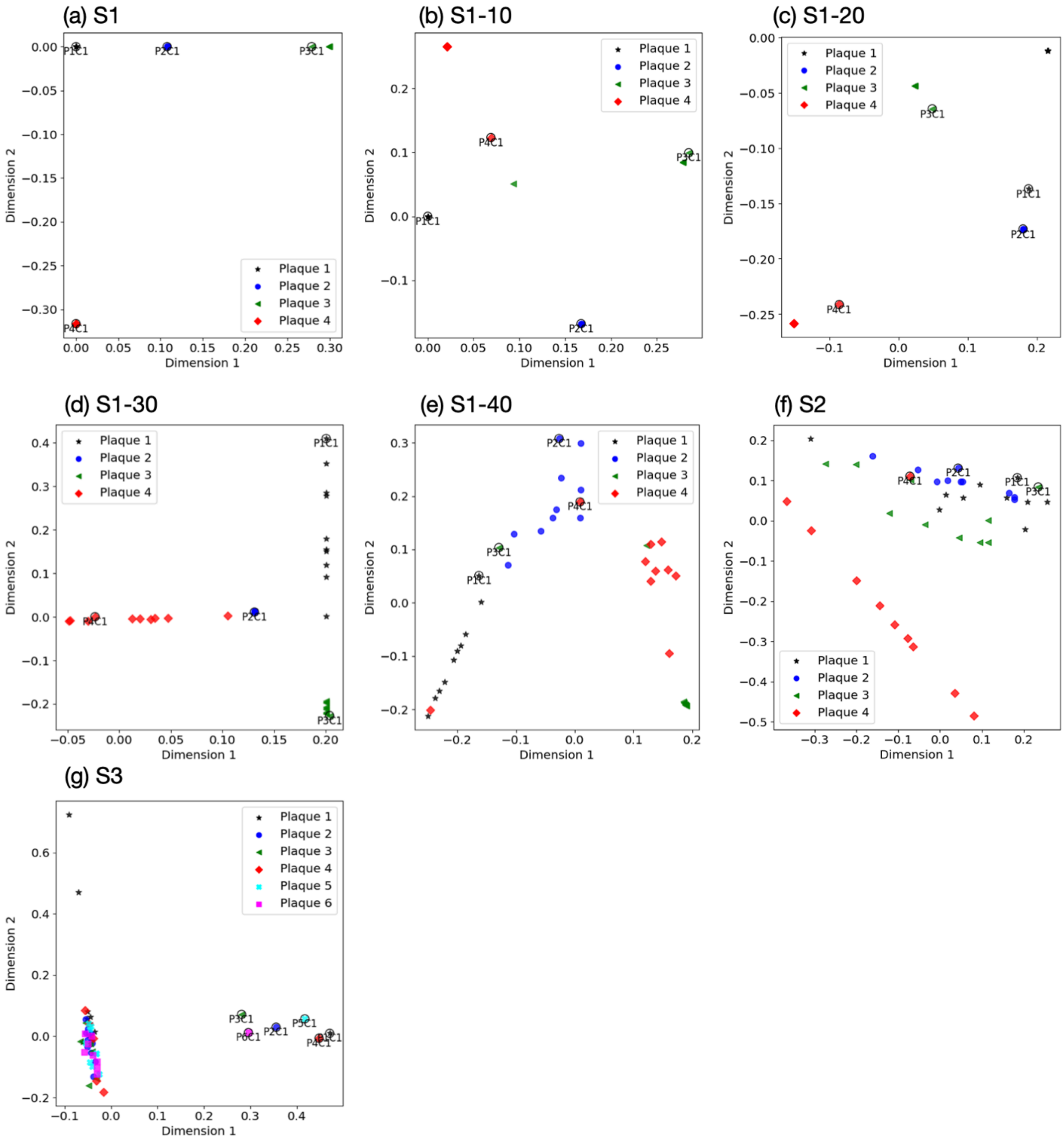


**Figure 13:** LLE analysis of toy datasets' attributes.

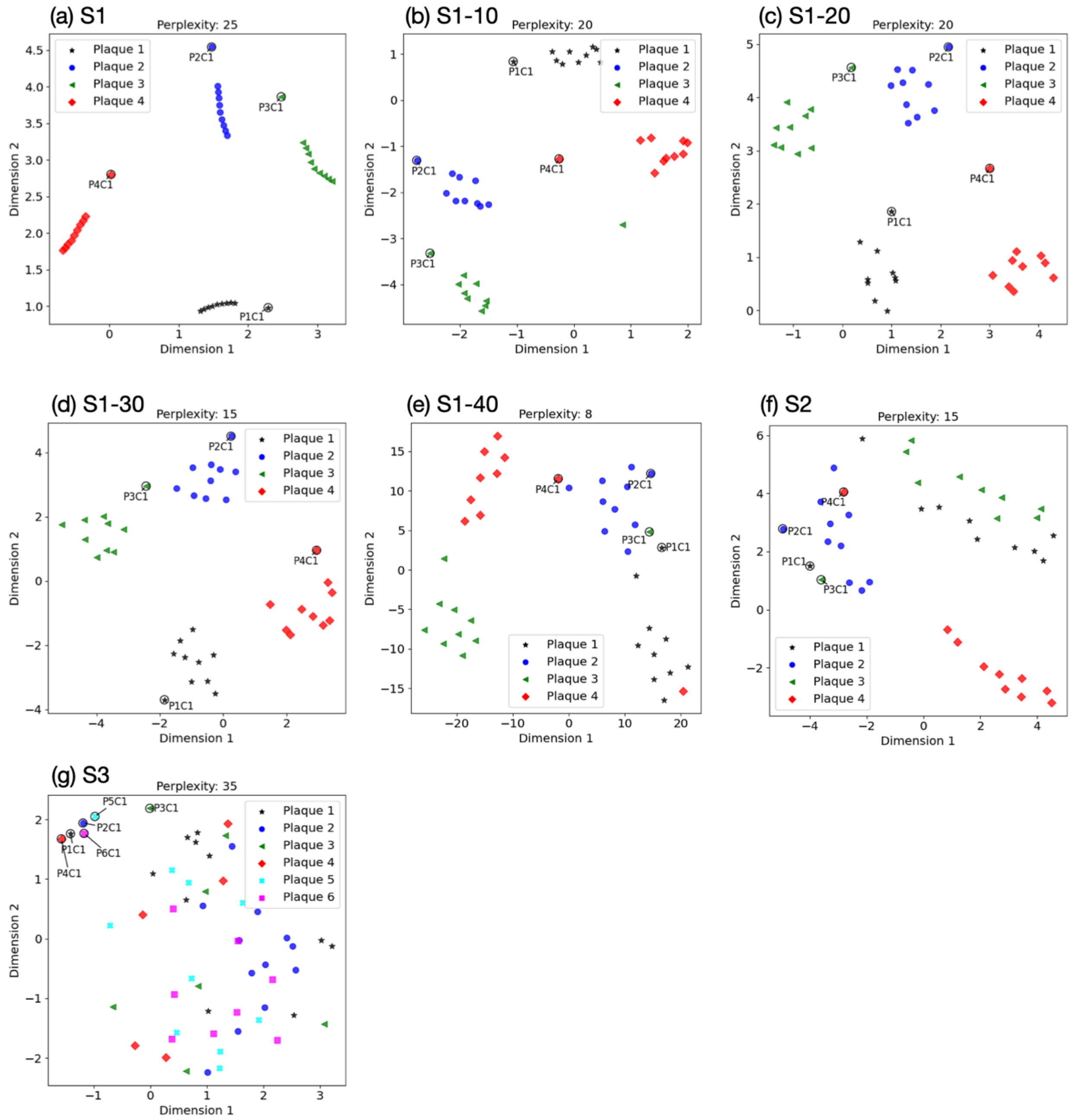


**Figure 14:** t-SNE analysis of attributes extracted from toy datasets

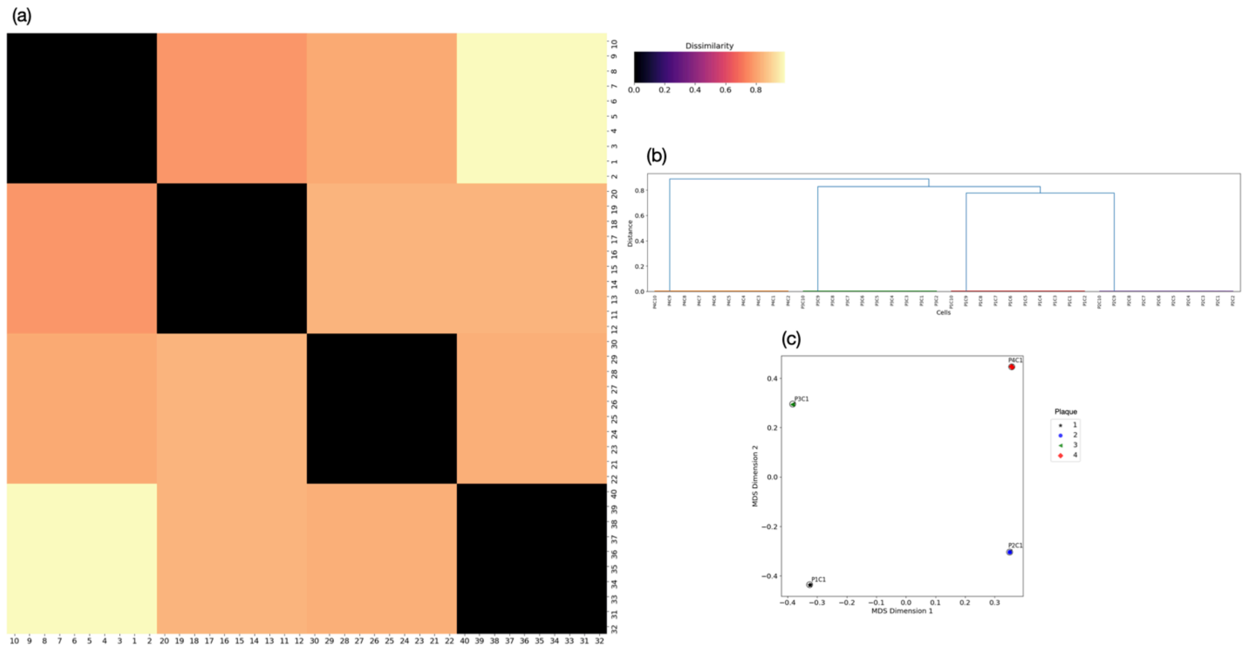


**Figure 15:** DTW analysis of S1: (a) heat map (b) dendrogram (c) two-component metric MDS of the dissimilarity matrix.

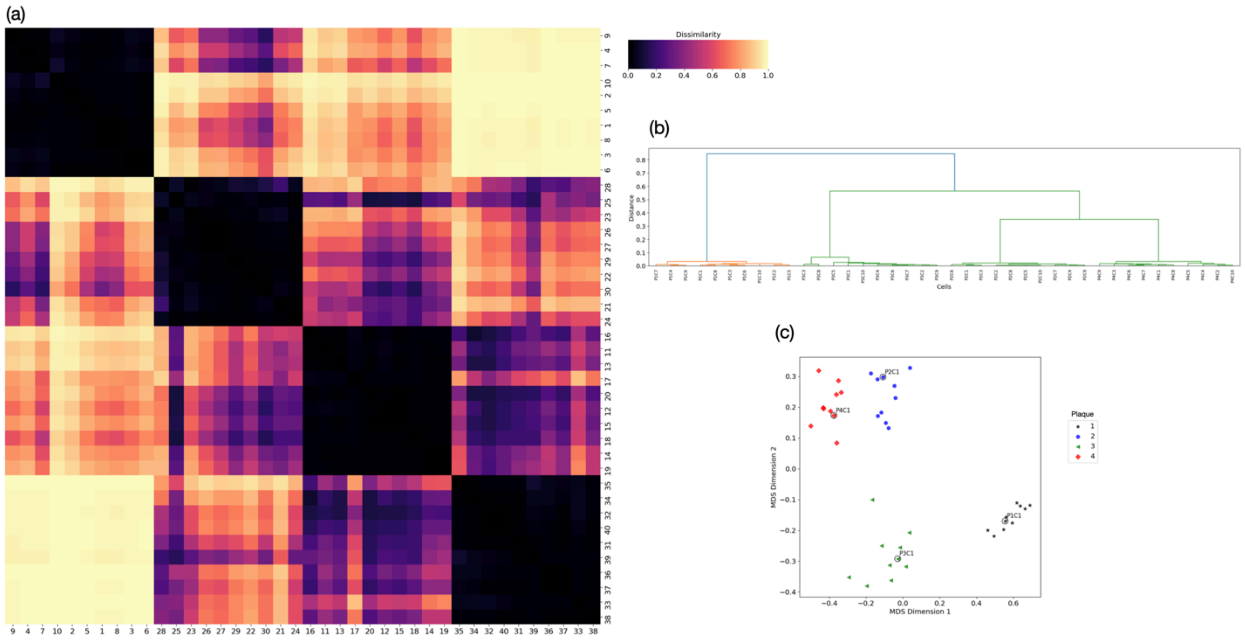


**Figure 16:** DTW analysis of S1-10: (a) heat map (b) dendrogram (c) two-component metric MDS of the dissimilarity matrix

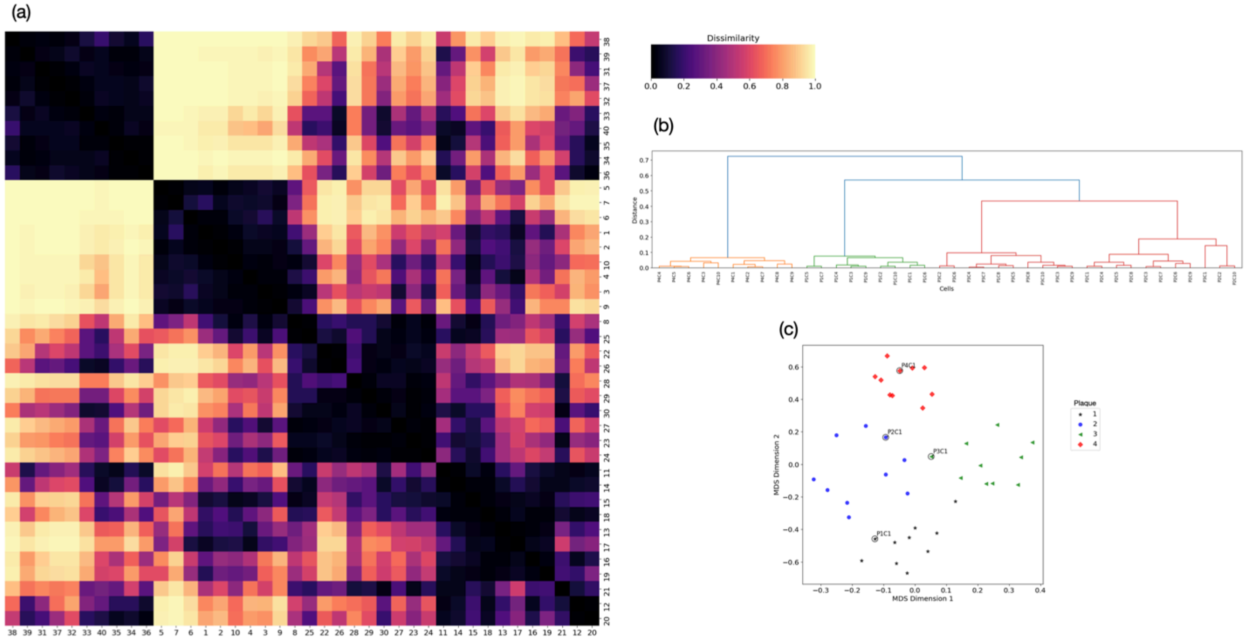


**Figure 17:** DTW analysis of S1-20: (a) heat map (b) dendrogram (c) two-component metric MDS of the dissimilarity matrix.

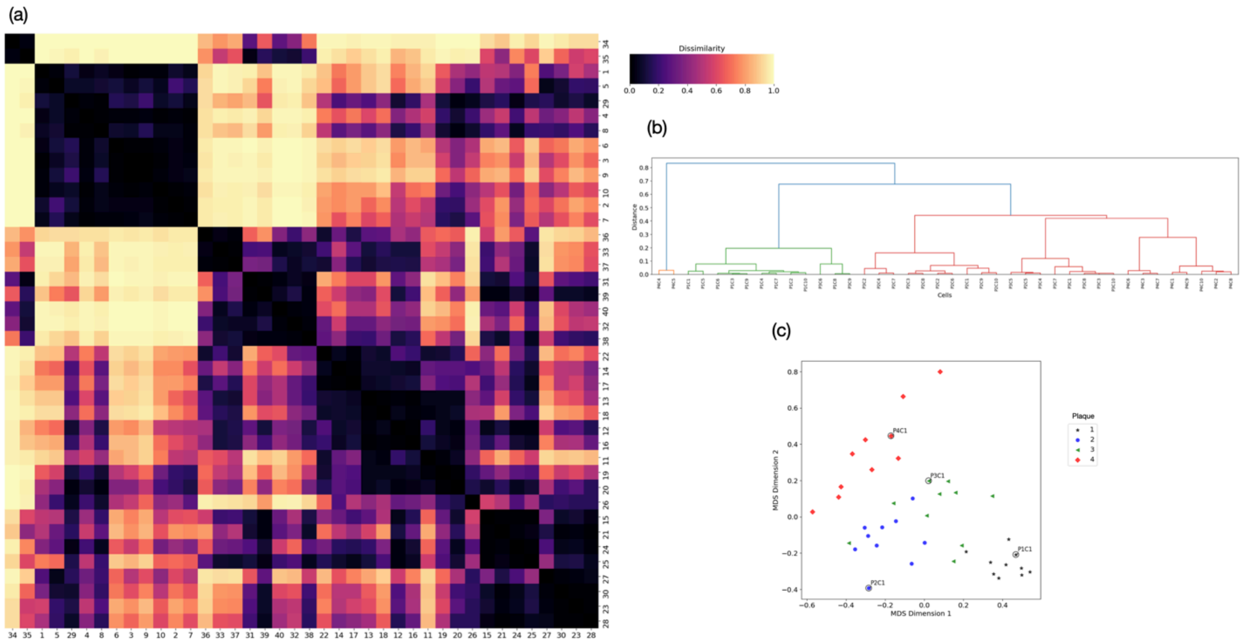


**Figure 18:** DTW analysis of S1-30: (a) heat map (b) dendrogram (c) two-component metric MDS of the dissimilarity matrix.

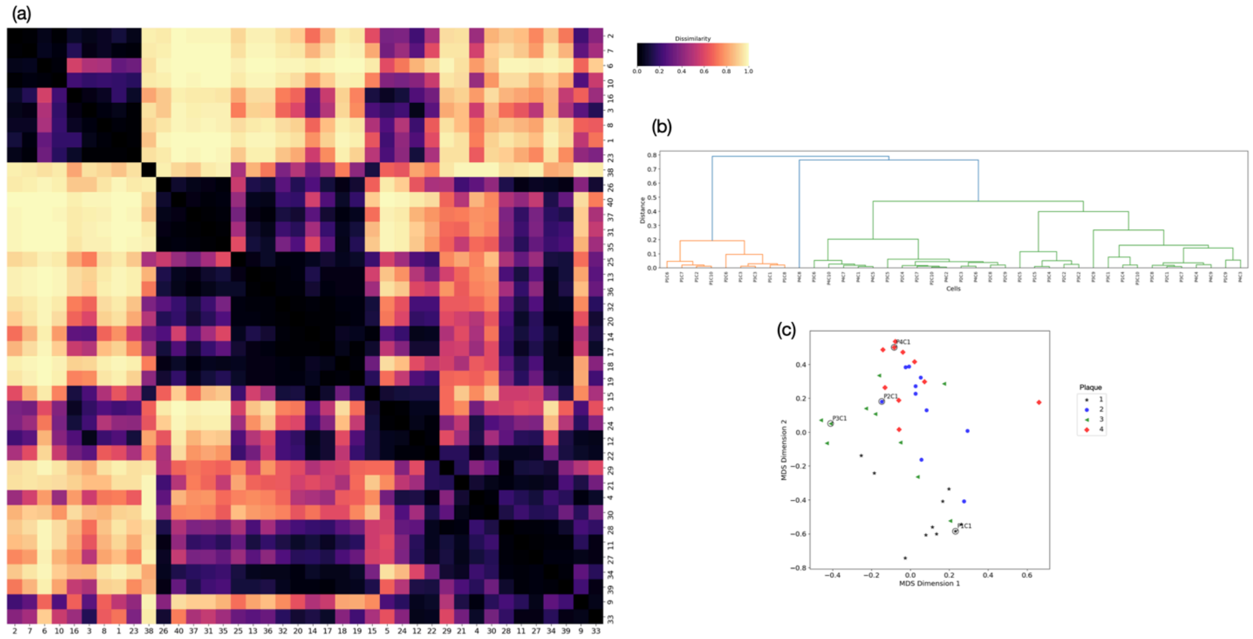

**Figure 19:** DTW analysis of S1-40: (a) heat map (b) dendrogram (c) two-component metric MDS of the dissimilarity matrix

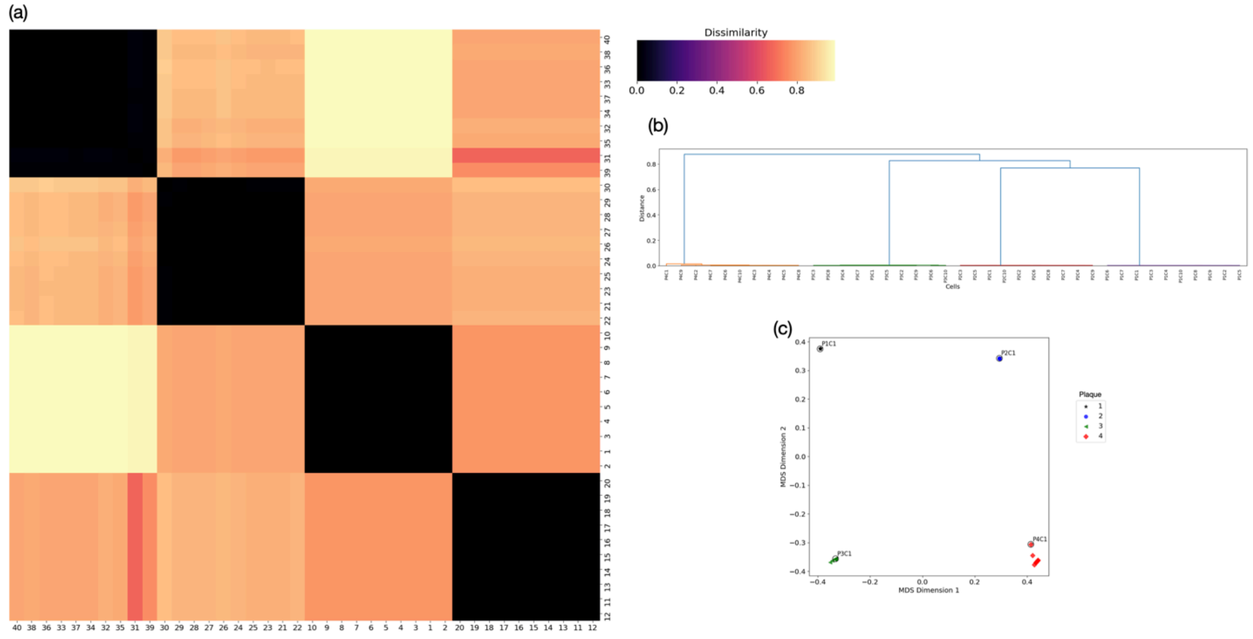


**Figure 20:** DTW analysis of S2: (a) heat map (b) dendrogram (c) two-component metric MDS of the dissimilarity matrix.

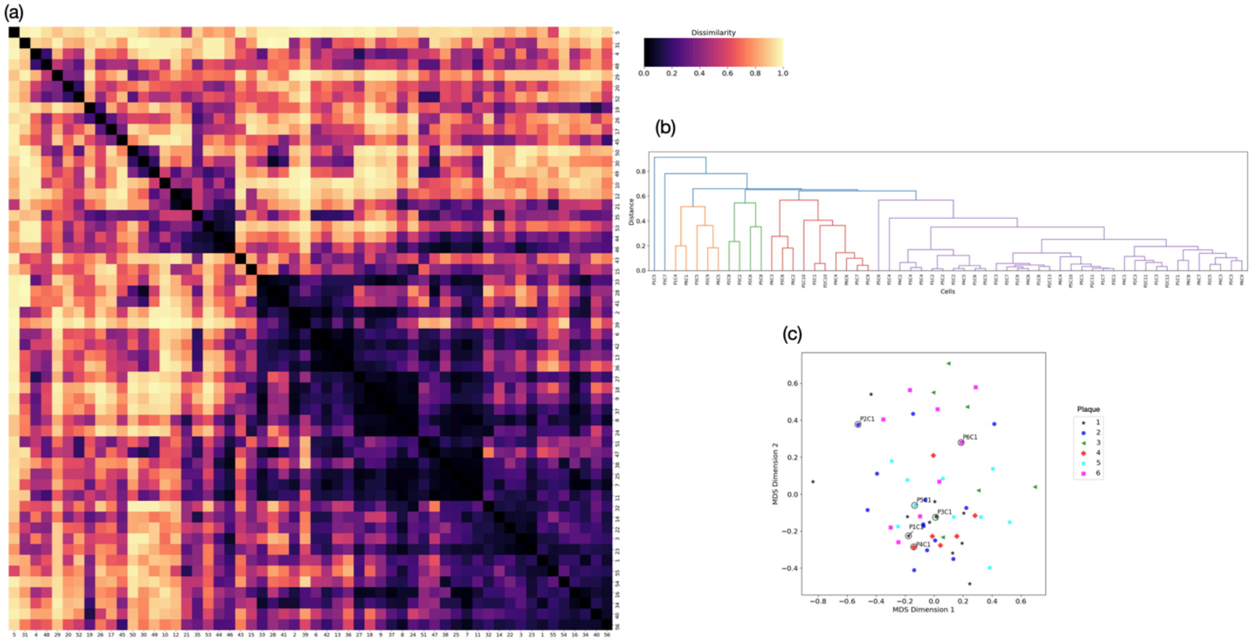


**Figure 21:** DTW analysis of S3: (a) heat map (b) dendrogram (c) two-component metric MDS of the dissimilarity matrix

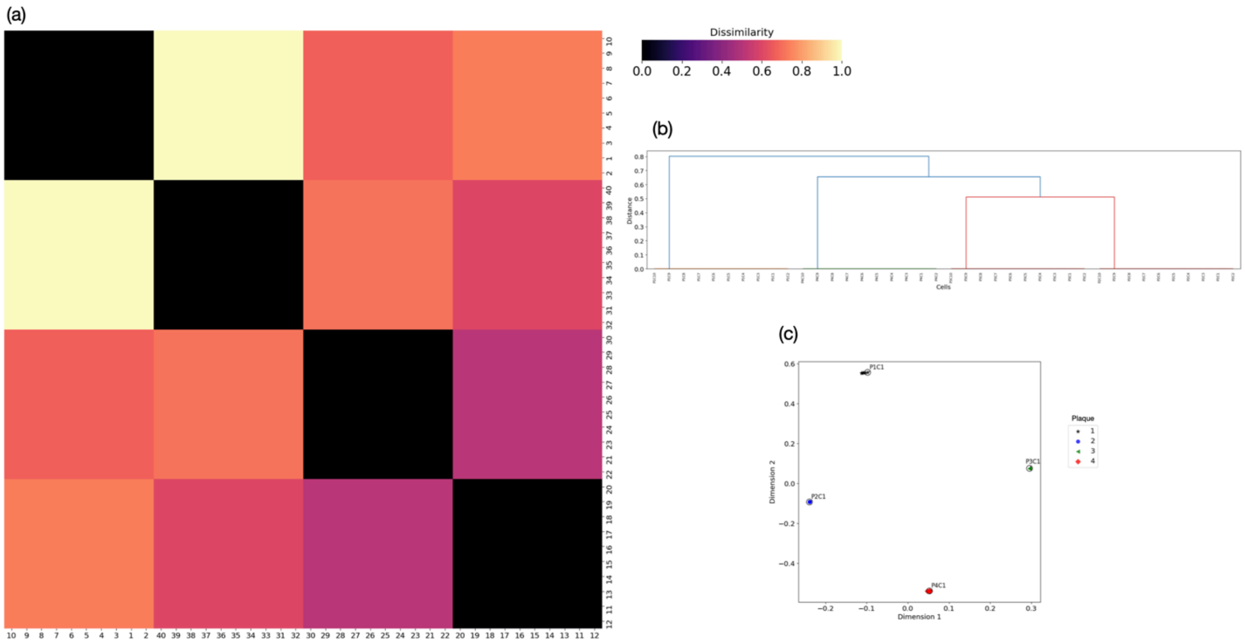


**Figure 22:** Analysis of S1 by RMSE method: (a) heat map (b) dendrogram (c) two-component metric MDS of the dissimilarity matrix

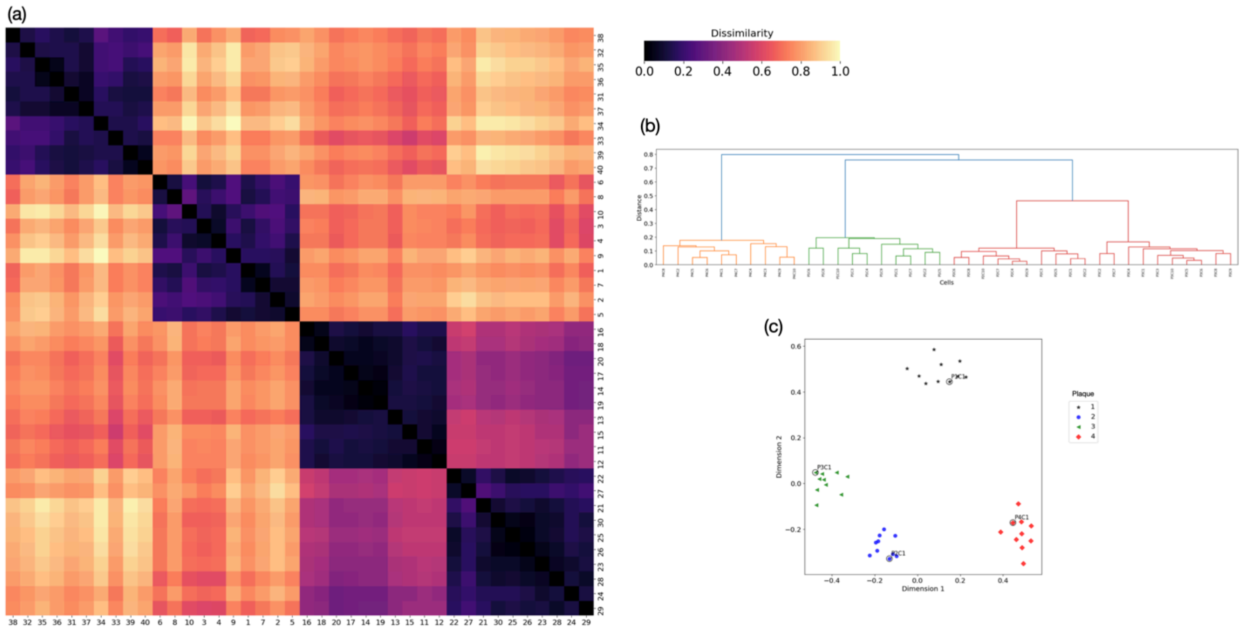


**Figure 23:** Analysis of S1-10 by RMSE method: (a) heat map (b) dendrogram (c) two-component metric MDS of the dissimilarity matrix

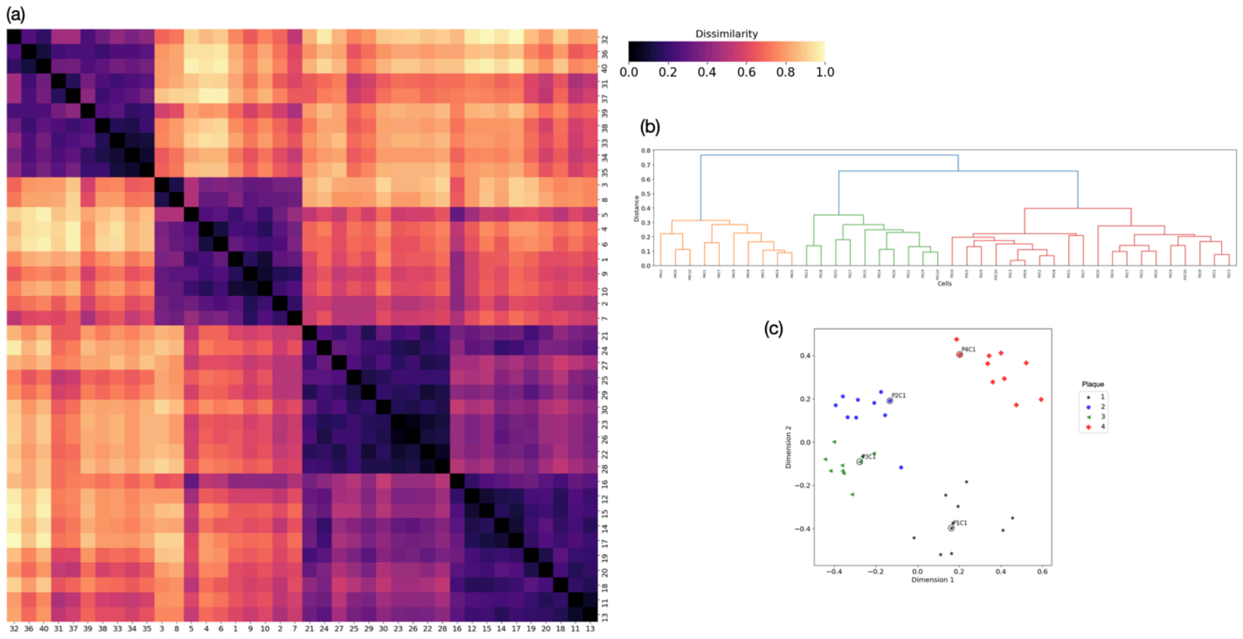


**Figure 24:** Analysis of S1-20 by RMSE method: (a) heat map (b) dendrogram (c) two-component metric MDS of the dissimilarity matrix.

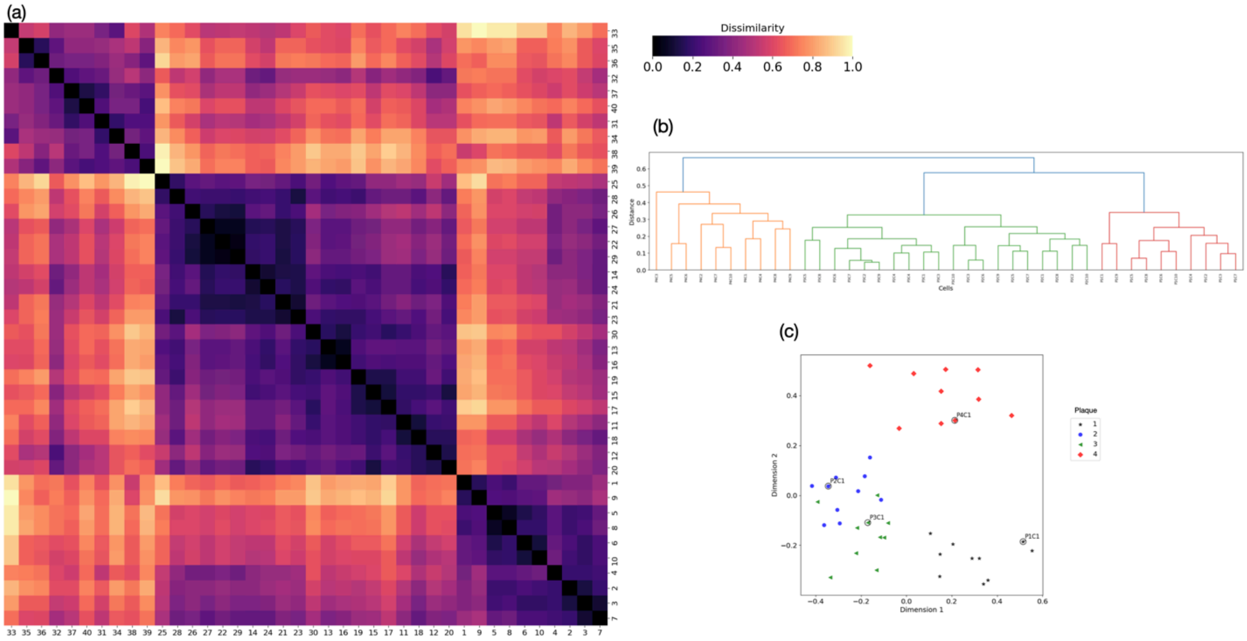


**Figure 25:** Analysis of S1-30 by RMSE method: (a) heat map (b) dendrogram (c) two-component metric MDS of the dissimilarity matrix.

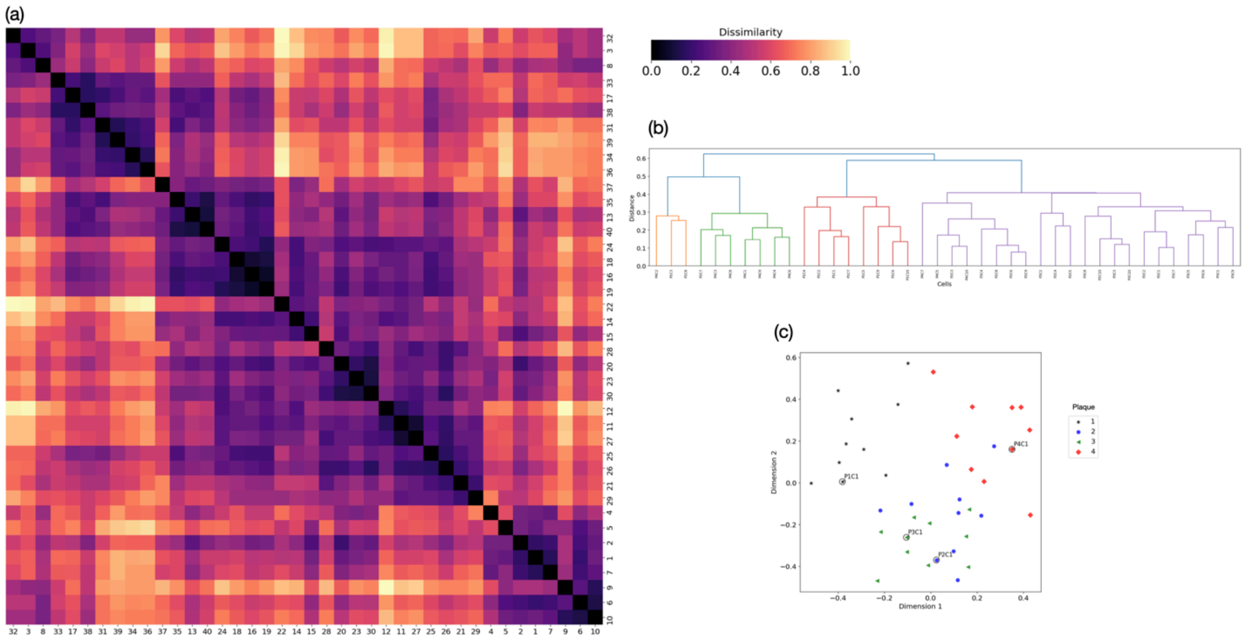


**Figure 26:** Analysis of S1-40 by RMSE method: (a) heat map (b) dendrogram (c) two-component metric MDS of the dissimilarity matrix.

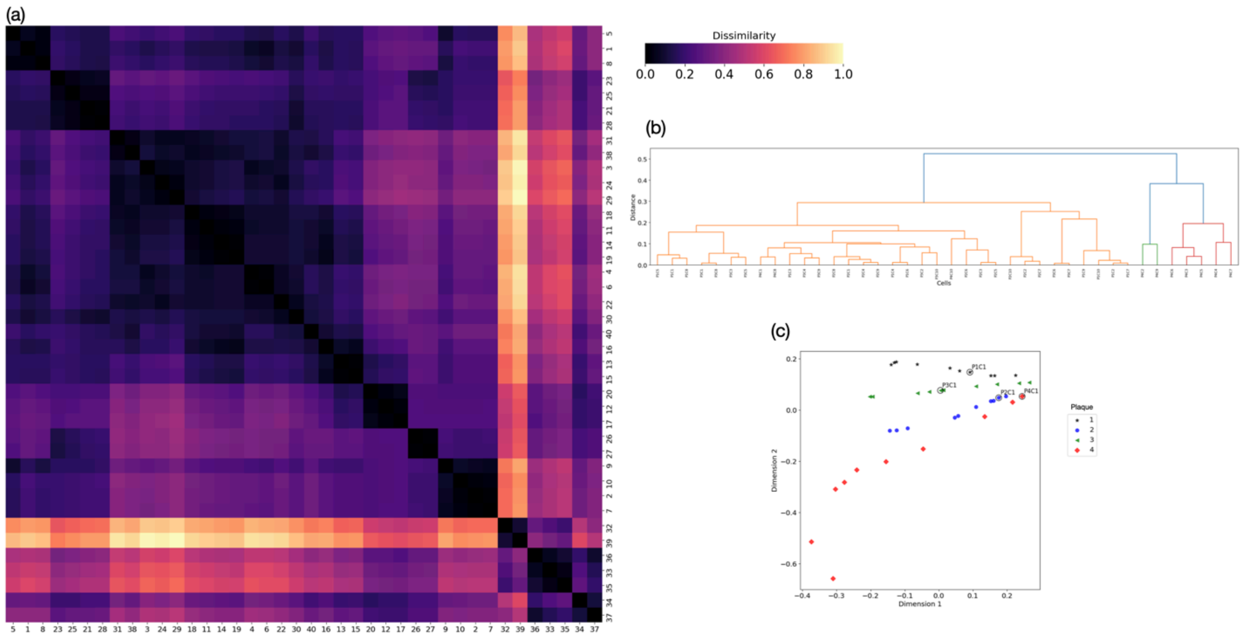


**Figure 27:** Analysis of S2 by RMSE method: (a) heat map (b) dendrogram (c) two-component metric MDS of the dissimilarity matrix.

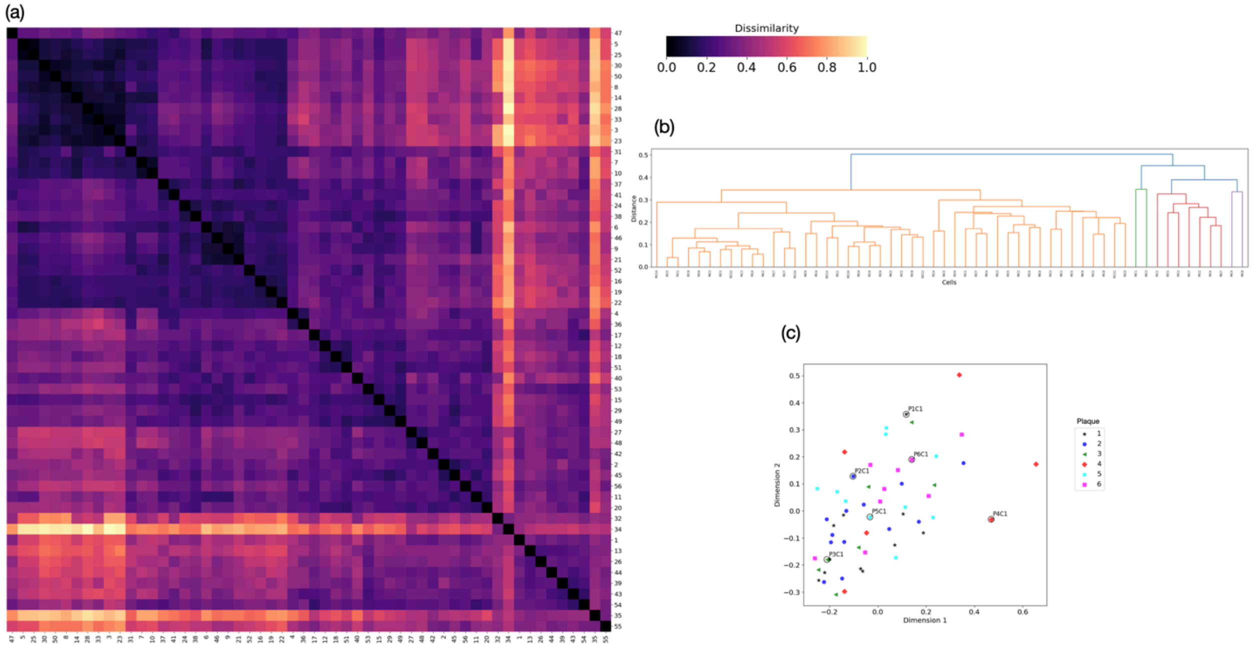


**Figure 28:** Analysis of S3 by RMSE method: (a) heat map (b) dendrogram (c) two-component metric MDS of the dissimilarity matrix.
